# Being a tree crop increases the odds of experiencing yield declines irrespective of pollinator dependence

**DOI:** 10.1101/2023.04.27.538617

**Authors:** Marcelo A. Aizen, Gabriela Gleiser, Thomas Kitzberger, Rubén Milla

## Abstract

Crop yields, *i.e.*, harvestable production per unit of cropland area, are in decline for a number of crops and regions, but the drivers of this process are poorly known. Global decreases in pollinator abundance and diversity have been proposed as a major driver of yield declines in crops that depend on animals, mostly bees, to produce fruits and seeds. Alternatively, widespread tree mortality has been directly and indirectly related to global climate change, which could also explain yield decreases in tree crops. As tree crops are expected to be more dependent on pollinators than other crop types, disentangling the relative influence of growth form and pollinator dependence is relevant to identify the ultimate factors driving yield declines. Yield decline, defined here as a negative average annual yearly change in yield from 1961 to 2020, was measured in 4270 time series, involving 136 crops and 163 countries and territories. About one-fourth of all time series showed declines in crop yield, a characteristic associated with both high pollinator dependence and a tree growth form. Because pollinator dependence and plant growth form were partially correlated, we disentangled the effect of each of these two predictors using a series of generalized linear mixed models that evaluated direct and indirect associations. Our analyses revealed a stronger association of yield decline with growth form than with pollinator dependence, a relationship that persisted after partialling out the effect of pollinator dependence. In particular, yield declines were more common among tree than herbaceous and shrub crops in all major regions but in Africa, a continent showing a high incidence of yield declines irrespective of growth form. These results suggest that pollinator decline is not the main reason behind crop productivity loss, but that other factors such as climate change could be already affecting crop yield.

## INTRODUCTION

Plant breeding has played a crucial role in improving agricultural productivity through techniques such as hybridization and polyploidization, artificial selection, and genetic engineering. Along with the expansion of agriculture and the intensification in the use of external subsidies (Matson and Vitousek 2006; Aizen et al. 2022), these methods have helped feed a growing human population (Borlaug 1983; Moose and Mumm 2008; Tester and Langridge 2010). However, the sustained increase in crop yields (*i.e.*, a positive change in harvestable production per unit of cropland area) of many crop species in different parts of the world is now decelerating, indicating that the increase in productivity might be reaching a ceiling (Bennett et al. 2012; Ray et al. 2012; Grassini et al. 2013). This may be in part because crop improvements are limited by various plant trade-offs that impose upper boundaries on yield growth (Huot et al. 2014; Fernandez et al. 2021; Garibaldi et al. 2021). Even though resource allocation patterns in crop plants may be driven to extremes by human selection, crops have rarely explored the phenotypic space existing beyond the boundaries set by the evolution of wild plants (Milla et al. 2018; Garibaldi et al. 2021; Cunha et al. 2023).

In addition to intrinsic constraints, environmental degradation associated with global change may cause declines in crop yield (*i.e.*, a negative change in harvestable production per unit of cropland area) over the last decades. In particular, the interaction of different plant traits with a plant’s changing abiotic and biotic environment may also contribute to diminishing yield growth, and result in yield decline (Bennett et al. 2012; Ray et al. 2012). One of the most influential factors affecting the yield of many crops is the availability of efficient pollinators, which have declined in many regions during the last decades due to a combination of habitat destruction, land-use change, intensive pesticide use, pathogen transmission, and climate change (Winfree et al. 2009; Potts et al. 2010; Zattara and Aizen 2021). Additionally, climate change has resulted in extreme temperature fluctuations, droughts, and flooding events (Coumou and Rahmstorf 2012; Stott 2016; Ummenhofer and Meehl 2017), which have increased plant stress (Greenwood et al. 2017; Onyekachi et al. 2019), pest susceptibility (Jaime et al. 2019; IPPC Secretariat 2021), and phenological mismatches (Beard et al. 2019). Two plant-related factors that can predict crop yield decline in relation to dwindling pollination services and climate change are pollinator dependence and growth form, respectively. Pollinator dependence can determine a crop’s susceptibility to changes in pollinator availability (Aizen et al. 2022), whereas a plant’s growth form and other correlated functional traits can determine susceptibility to extreme weather events and plasticity to respond to global warming (Greenwood et al. 2017; Alecrim et al. 2023).

Dependency on pollinators varies greatly among crops (Aizen et al. 2022). On one extreme, some crops grown for their vegetative organs (*e.g.*, potato, carrot, tea, etc.) or their wind-pollinated seeds or fruits (*e.g.*, wheat, maize, olive, etc.) do not rely on pollinators to produce the parts we consume. On the other extreme, some seed and fruit crops have a high degree of pollinator dependence (*e.g.*, cacao, watermelon, vanilla, etc.), to the point that their yield would be reduced close to zero in the absence of pollinators (Klein et al. 2007). However, more than half of all cultivated crops fall somewhere between these two extremes, which means that their yield can be improved to different degrees in the presence of pollinators (Aizen et al. 2009, 2022). In any event, the presence of a diverse group of pollinators, sometimes including rare but highly effective pollinators, is relevant for increasing the yield of most pollinator-dependent crops, particularly of those with high dependency (Garibaldi et al. 2013; Sáez et al. 2022). For instance, the yield of several crops tends to decrease with increasing distance from the field edge of the cultivated field in association with a decline in the abundance and diversity of wild pollinators that thrive in field margins (Garibaldi et al. 2011b). In addition, the yield of economically important tropical crops such as coffee, which benefit from the pollination provided by diverse pollinator assemblages, has declined in different countries (Aizen et al. 2020). A recent meta-analysis revealed that relying solely on honey bees (*Apis mellifera*), the most important managed pollinator globally, is insufficient to significantly reduce the pollination deficit of most cross-pollinated crops (Sáez et al. 2022). Based on these findings, it is reasonable to expect that the degree of a crop’s dependence on pollinators will influence the yield response to declines in pollinator populations. Specifically, we predict that as the dependency on pollinators increases, the occurrence of negative trends in crop yield over the last few decades is expected to increase as well.

The growth form of plants can determine their susceptibility to global warming. Trees are expected to show much less adaptive plasticity than herbs to a rapidly changing climate because of their longer lifespans and slower growth rates, with shrubs characterized by intermediate life-history attributes. For instance, spring-flowering forest herbs are advancing their phenologies faster than trees, thus taking advantage of the longer growing season (Alecrim et al. 2023). On the other hand, trees have deeper root systems and greater leaf biomass than shrubs and herbs, which allows them to tap water and nutrients from deeper in the soil and capture more sunlight for photosynthesis. However, these traits could also make trees more vulnerable to changes in temperature and precipitation because, in addition to their more stringent hydraulic limitations (Choat et al. 2012), they require more resources to support their growth and metabolism. In particular, secondary growth in trees and shrubs requires a significant amount of energy and resources to produce the lignin that makes up the tree’s woody tissue (Novaes et al. 2010), which could increase their susceptibility to stress factors, such as drought and pests (Cailleret et al. 2017). Also, increasing occurrences of wildfires can cause direct damage to the cambium layer responsible for wood formation (Dickinson et al. 2004), whereas extreme temperatures and increasing incidence of droughts and frosts can cause xylem embolism and cavitation (Martínez-Vilalta and Pockman 2002; Savi et al. 2015), all leading to growth abnormalities, partial crown dieback, and increased mortality (Barigah et al. 2013; Cailleret et al. 2017; Greenwood et al. 2017). On the other hand, herbaceous plants can experience reduced reproduction and high mortality in relation to extreme temperatures; however, they are probably better adapted to overcome unsuitable climatic episodes as they can survive as dormant seeds or underground structures (Keeley et al. 1981; Gardarin and Colbach 2015; Jongen et al. 2015). In an agricultural context, long-lasting yield declines in tree crops can be triggered by sporadic but increasingly frequent heat waves, frost events, or pest outbreaks. On the other hand, extreme climatic events or pest outbreaks do not have long-lasting consequences in herbaceous crops, most of them annuals, as, unlike long-lived crops, they are sown anew every year and breeding can provide adaptations within a shorter time frame. Given this background, we expect tree crops to exhibit a higher incidence of long-term yield declines than herb crops, whereas shrub crops would fall in between.

The effects of pollinator dependence and growth form on yield decline cannot be studied independently because these two factors are expected to be associated for two reasons. First, most crops harvested for their vegetative parts, which are thus pollinator-independent, are herbs (Klein et al. 2007). Second, among crops cultivated for their fruits and seeds, pollinator dependence is expected to increase from herbs to trees because the incidence of self-incompatibility, which implies mandatory cross-pollination for successful fertilization, is higher in long-lived plants (Ramírez 2022; Cunha and Aizen 2023). Thus, the effect of pollinator dependence on the probability of yield decline can be easily confounded with the effect of growth form and vice-versa when both factors are considered separately.

Here we investigated the relationship between pollinator dependence and growth form across crops and then analyzed whether either pollinator dependence or growth form is associated with the probability of crop yield decline over the last six decades. We examined the effects of pollinator dependence and growth form separately and jointly after accounting for any confounding effect associated with total cultivated area per crop, which might also relate to yield decline (Aizen et al. 2009). Although our assessment is global, whether a crop’s yield declines or not was estimated at the country scale (the smallest spatial scale available for long-term data), rather than at the regional or global scale. This relatively small scale was chosen because declines in yield in some countries may be compensated by increases in others, thus hiding any effect that spatially heterogeneous either abiotic, biotic, economic or political factors may have on yield decline. Compilation and analysis of country-level data also allow for the detection of geographical areas where yield decline may be more prevalent, and to determine whether potential associations between the probability of yield decline and either pollinator dependence or growth form vary among regions or can be regarded as a truly global trend.

## MATERIALS AND METHODS

### The database

We obtained yearly data (1961-2020) on yield and cultivated area at the country level from the United Nations Food and Agriculture Organization database (FAOSTAT 2021) for a total of 136 crops and crop items (*i.e.*, aggregations of different species or subdivisions in terms of different harvested parts; see Aizen et al. 2019 for details) for which there is available information on pollinator dependence (Klein et al. 2007; updated and expanded in Aizen et al. 2022) and there are uninterrupted yield records since 1961 for at least one country. Data were retrieved for 163 countries and territories that maintained their physical integrity since 1961, resulting in a total of 4280 complete 60-yr time series (crop x country combinations). We discarded 10 series that showed unrealistic differences in yield (> two orders of magnitude) between any two years.

For classifying crops into pollinator-dependence categories, we considered initially the five categories established by Klein et al. (2007) based on the expected reduction in crop yield in the absence of animal pollination: none (0% reduction), little (>0 to <10%), modest (≥10 to <40%), high (≥40 to <90%) and essential (≥90%). However, given the highly unbalanced number of crops in each category (see Table S1), particularly when pollinator dependence was crossed with growth form, we reduced the number of categories to three by maintaining the category “none” and merging the categories “little” and “modest” into the category “modest” and the categories “high” and “essential” into the category “high”. Among pollinator-independent crops (category “none”), we further distinguished among those cultivated for their vegetative parts (*i.e.*, leaves, stems, bark, roots, tubers, etc.) *vs.* reproductive parts (*i.e.*, either fruits or seeds). By definition, all pollinator-dependent crops (our categories “moderate” and “high”) were cultivated for either their fruits or seeds. Finally, crops were classified into one of three commonly used growth-form categories (*i.e.*, herbs, shrubs, and trees) using existing databases (Milla 2020; Gleiser et al. 2021). As a result, the two main focal factors, pollinator dependence and growth form, had the same number of categories and degrees of freedom, which make them comparable in terms of inferential testing and associated statistical power (Cottingham et al. 2005). To explore for potential geographical differences in yield decline associated with either pollinator dependence or growth form, countries were grouped into one of the five following geographical regions defined by the United Nations Geoscheme (https://unstats.un.org/unsd/methodology/m49/): Africa, the Americas (including South, Central, and North America), Asia, Europe, and Oceania (including Australia and New Zealand).

### Data analyses

For each time series, we estimated the yield growth rate (*i.e.*, the average annual yearly change in yield) as the slope of the least-squares linear regression of (log) yield *vs*. year (Fig. S1). This simple statistical method provides similar point estimates as other regression methods that consider the autocorrelation nature of time series (Altinay 2003). Given the goal of our study, an average growth rate <0 over the period 1961-2020 was considered evidence of yield decline, independent of the absolute value of the growth rate and without imposing any extra arbitrary criteria or value thresholds. This simple dichotomous classification provided clear evidence of whether the yield a given crop in a given country was declining or not for the vast majority of trends (see Results).

In terms of statistical analyses, we first evaluated the association between pollinator dependence and growth form by means of a chi-square test. Then, we ran a step-up, hypothesis-driven series of sequential generalized linear mixed-effects models (GLMMs) to disentangle the influence of pollinator dependence and growth form on yield decline, considering the response variable as dichotomous based on the sign of the yield growth rate (negative slope=declining yield, coded as 1; positive slope = non-declining yield, coded as 0). The first model (model GLMM_0) was a pure random model to characterize variation among crops (independent of country) and countries (independent of crop) in yield decline, and thus only included crop and country as random crossed factors. This model was aimed at obtaining “raw” descriptive estimates of yield decline by crop and country. In addition to the random factors crop and country, the second and third models included pollinator dependence (GLMM_1a) and growth form (GLMM_1b), respectively, while the fourth model (GLMM_2) included both factors together to assess the effect of pollinator dependence independent of growth form, and vice versa. GLMM_1 and GLMM_2 also included the geographical region as another fixed factor to account for regional variation in yield decline and, more specifically, to test for differences in yield decline between regions. A last model (GLMM_3) assessed whether the effect of pollinator dependence depended on growth form (i.e., the “pollinator dependence x growth form” interaction), and whether the influence of either of these two crop factors on yield decline depended on geographical region (i.e., the “pollinator dependence x region” and “growth form x region” interactions). All the mixed models *sensu stricto* (*i.e.*, GLMM_1a, GLMM_1b, GLMM_2, and GLMM_3) included the (log10) cumulative total harvested area (in square kilometers) over the period 1961-2020 for each crop (summed across countries) and for each country (summed across crops) as covariates to account for variation among crops and among countries in the probability of yield decline that could relate to the agricultural relevance of the crop and country, respectively. The log-transformation of these two variables was necessary because the raw data encompass about six orders of magnitude with an extremely right-skewed distribution. Using the same random structure and area covariates as in the former models but considering only data for pollinator-independent crops, we tested for differences in the probability of yield decline between crops cultivated for their reproductive *vs*. vegetative organs.

All models above considered a binomial (0,1) distribution and a *logit* link function and were implemented with the statistical software R version 4.0.2 (R_Core_Team 2020), using the glmmTMB function of the glmmTMB package (Brooks et al. 2017) and rechecked for consistency using the glmer function of the lme4 package (Bates et al. 2015). Because both functions prouced highly similar results (only differing at the second or third decimal in their parameter estimates), we only report here results from the models run with glmmTMB. The extent of multicollinearity among all fixed factors and covariables was assessed using the adjusted Generalized Variance Inflation Factor (GVIF, adjusted GVIF^[1/(2*df)], where df stands for degrees of freedom). GVIFs were estimated using the vif function from the car package (Fox and Weisberg 2019), where adjusted GVIF values <3 indicate no serious multicollinearity problems. Effects were evaluated statistically using Wald’s type II tests because we were firstly interested in assessing the overall average effects and secondarily the effects of the interactions. Finally, pairwise means were compared using Tukey’s test for multiple comparisons by running the contrast function of R’s lsmeans package (Lenth 2016).

Ignorance of phylogenetic non-independence due to shared evolutionary history among crops could bias results and inflate type-I errors. However, no need for a phylogenetic correction, with the consequent loss of statistical power, is required when phylogenetic regression models do not outperform equivalent non-phylogenetic regression models and there is no evidence of phylogenetic structure in the residuals of the non-phylogenetic models (Revell 2010). To evaluate these two conditions, we first constructed a phylogeny for the 136 crops following the protocol of Gleiser et al. (2021) and Milla (2020). In brief, obtaining the crop phylogeny involved checking accepted scientific species names based on the Plant List (The_Plant_List 2013) and pruning the GBOTB.extended megaphylogeny (Jin and Qian 2019) according to the unique scientific names assigned to each of the 136 crops. In the case of crops including a taxonomically diverse group of species, the whole group was assigned to the most representative species within the group, provided that the vast majority of groups are composed of a single species that is agriculturally relevant plus other minor crops (Milla and Osborne 2021). Then, we compared all the above non-phylogenetic models with homologous generalized linear mixed linear models with the same fixed and random structures, but that also included a correlation matrix based on the phylogenetic distances between crops (Revell 2010). These models were run using the phylo_glmm function written by Li and Bolker (2019) in R language (R_Core_Team 2020). Rather than imposing a given phylogenetic correlation structure on the random effects, this function models trait evolution following a flexible Brownian motion process that in practice is implemented as a sequence of independent errors (Li and Bolker 2019). Corresponding models with and without phylogeny were compared using Akaike Information Criterion (AIC). Also, residuals of phylogeny-ignorant models were averaged by crop species and these averaged residuals were analyzed for evidence of a phylogenetic signal using Blomberg’s *K* (Blomberg et al. 2003) estimated by using the R’s package phytools (Revell 2012).

## RESULTS

### The incidence and distribution of yield decline

We found that 22.58% (n=964) of the 4270 times-series compiled and analyzed were characterized by negative growth rates in yield over the period 1961-2020. Consistent with the proposal that a negative growth rate can be taken as evidence of long-term yield decline, the frequency distribution of the year of maximum yield across the 964 time-series exhibiting negative growth rates peaked in 1961, the first year of the 1961-2020 time series. About 90% of all these time series had a reduction of >5%, and 82.5% a reduction of >10% in yield over the whole period. In contrast, the frequency distribution of the year of maximum yield across the 3306 time-series with positive growth rates peaked in 2020, the last year of the 1961-2020 time series (Fig. 1).

**Fig. 1.**
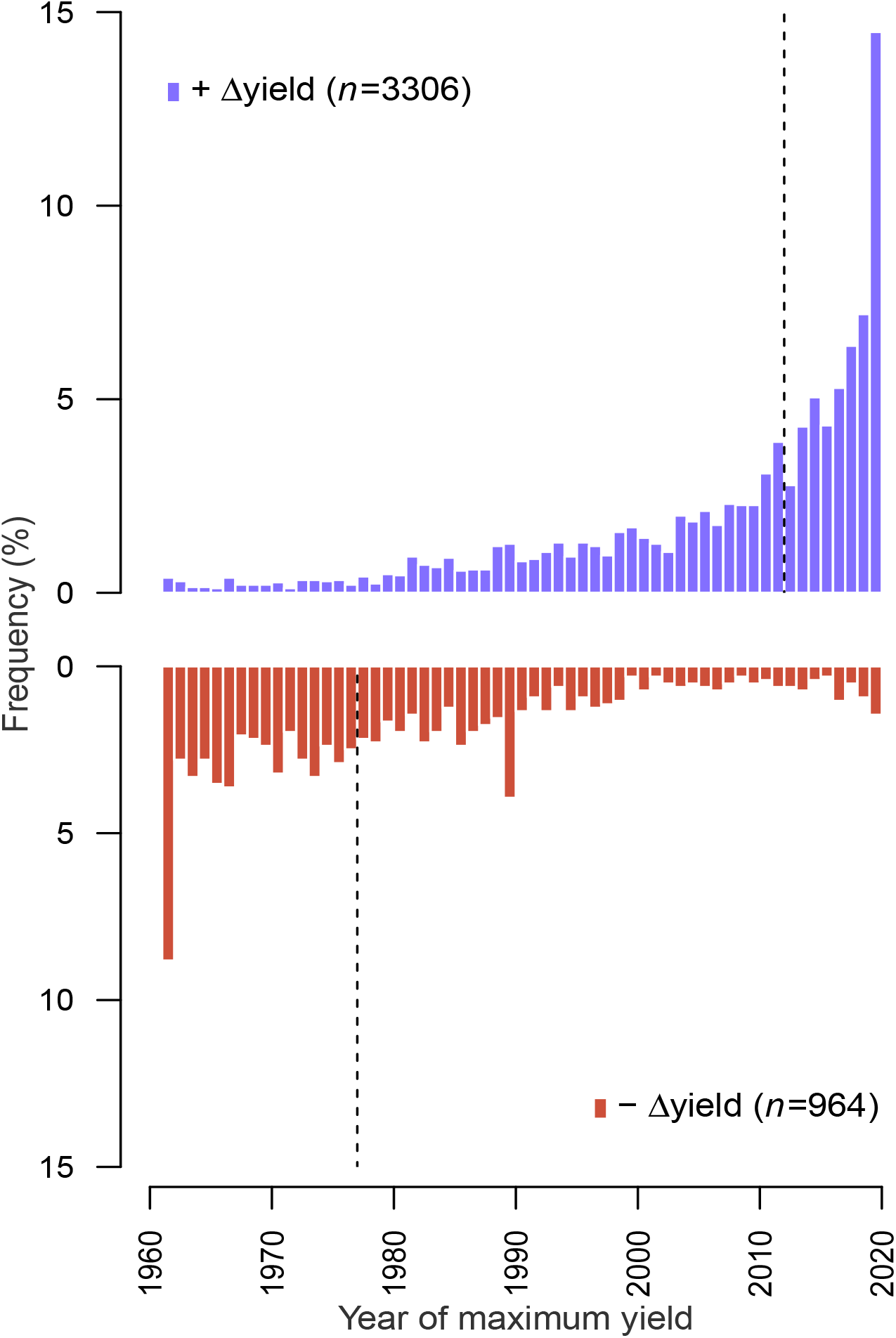
Relative frequency of year of maximum yield for 4270 crop x country yield trends (1961-2020) categorized as trends showing positive or negative average yield growths. The dashed lines indicate the median year of maximum yield for each yield-growth category.

The expected mean probability of yield decline estimated by the random model (GLMM_0) was 0.234 (95%CI=[0.215, 0.255]). However, there was high variability in the incidence of yield decline among crops and countries. The expected probability of yield decline varied from <0.1 in widespread crops such as maize, wheat, and rape, to >0.4 for assorted crops such as dates, cherries, walnuts, and cauliflowers (Fig. S2). The number of crops per country ranged from one in small island states like Nauru and Tuvalu to 76 in Turkey, with a median of 23 crops (Table S2). Variation among countries in the probability of yield decline was even more diverse than among crops, ranging from <0.08 in countries such as Hungary, Turkey, Myanmar, Turkey, and China, to >0.50 in countries such as Ecuador, Mauritius, Zimbabwe, and Trinidad and Tobago (Fig. S3). Particularly, there seems to be a high concentration of countries with a high probability of yield decline in Central and Southern Africa, Oceania, and to a lesser extent, the Pacific rim of South America (Fig. 2). Regional differences in the probability of decline were strong (Table 1), with countries in Africa and Oceania, followed by countries in the Americas, depicting the highest expected probability of yield decline (Fig. S4).

**Fig. 2.**
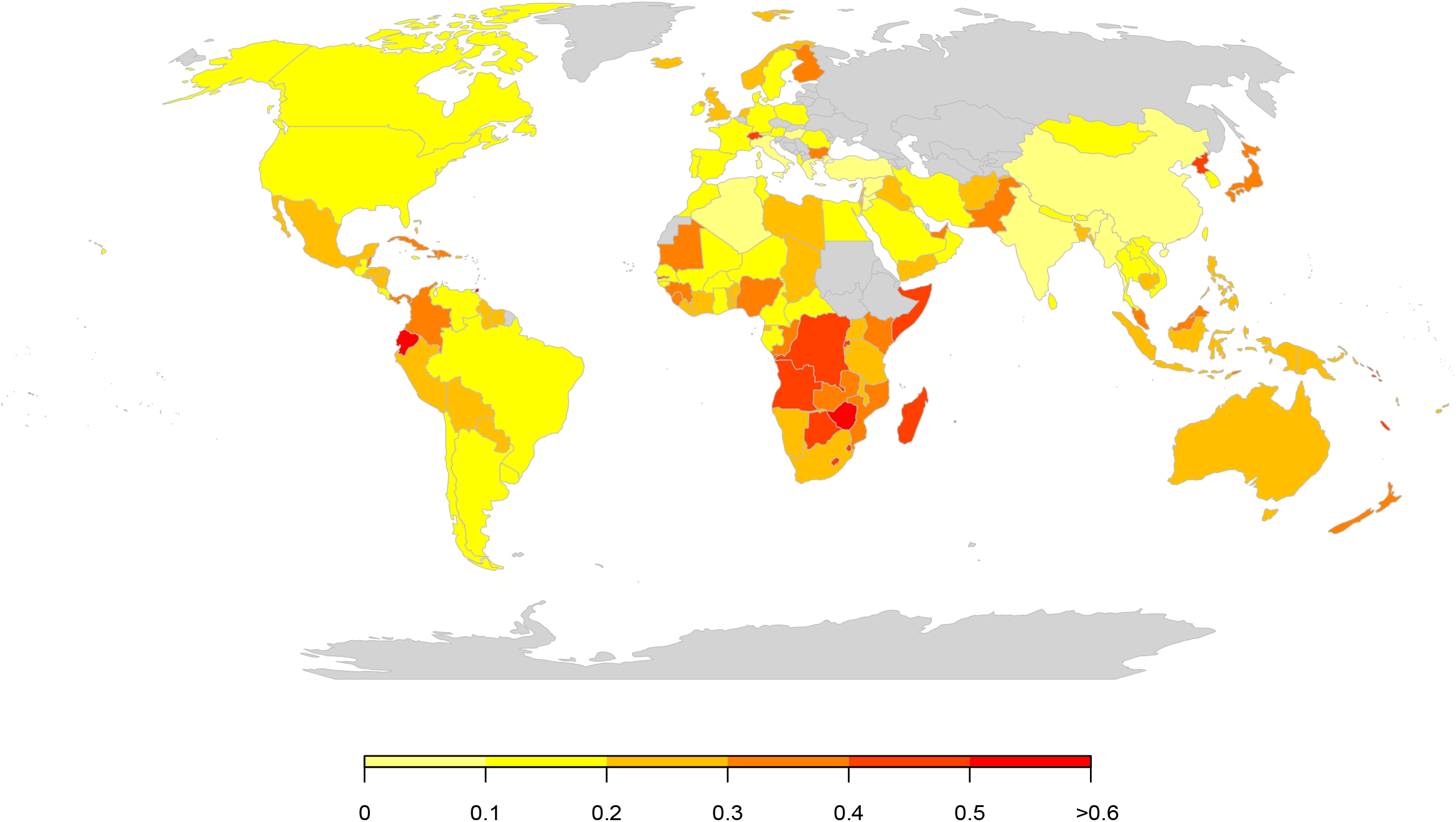
World map of yield decline. The map depicts the probability of yield decline (*i.e.*, the estimated proportion of crops that showed negative average growth rates in each country from 1961-2020) according to model GLMM_0 (see also Fig. S3).

**Table 1.**
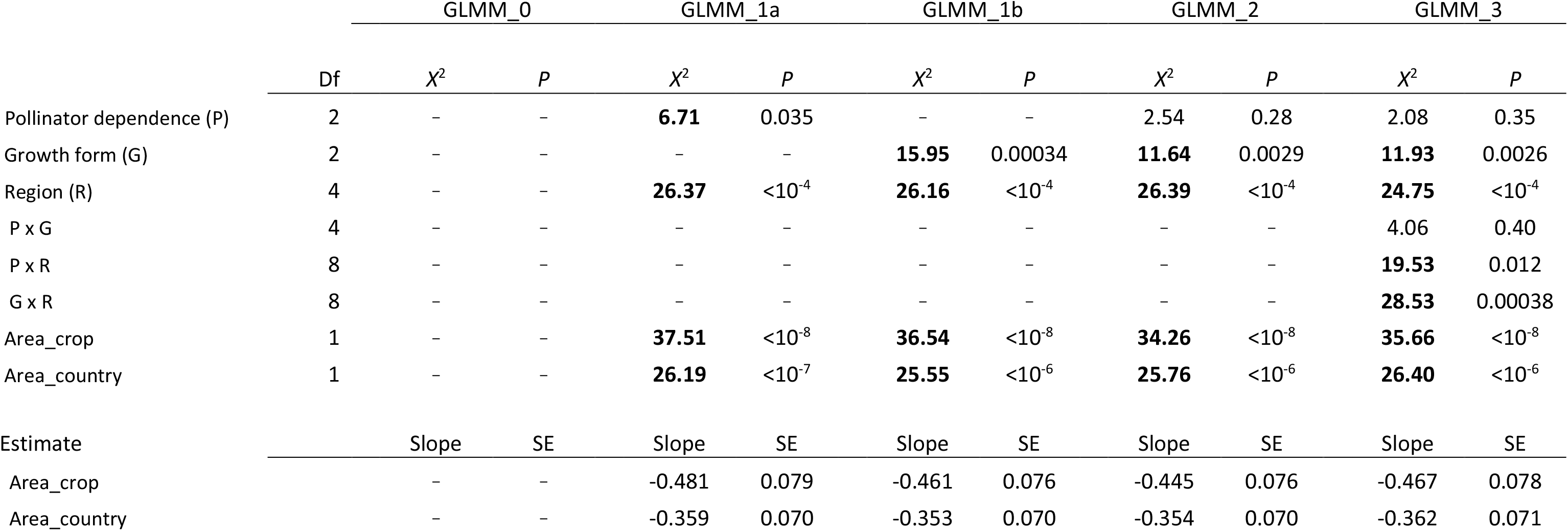
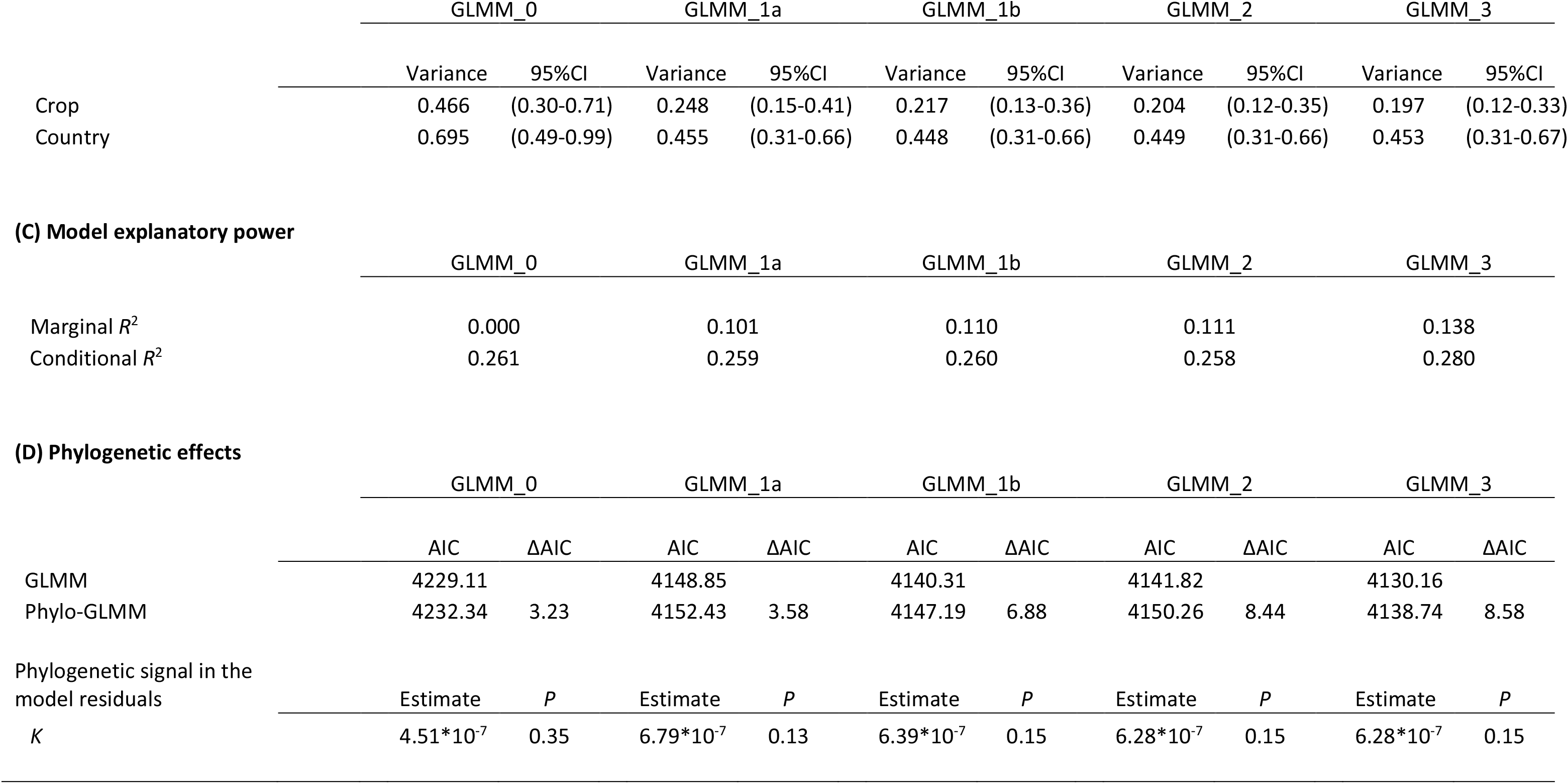
Results of logistic mixed-model analysis assessing the effects of pollinator dependence, growth form, and geographical region on the probability of yield decline (*i.e.*, the probability that a given crop in a given country shows an average annual growth rate in yield <0 over the period 1961-2020). GLMM_0 only evaluates the extent of (random) variation in the probability of yield decline among crops and countries. All the other four models include the effects of geographical region, and (cumulative) cultivated area per crop and country as fixed effects. Model GLMM_1a and model GLMM_1b test whether variation among crops in the probability of yield decline can be accounted for by pollinator dependence or by growth form, respectively. Model GLMM_2 tests for potential confounding effects as both factors, pollinator dependence and growth form, are associated to some extent (Fig. 3). Last, model GLMM_3 assesses whether any effect of pollinator dependence depends on growth form, or whether any effect of crop pollinator dependence or growth form differs among regions. Fixed effects (A) are evaluated statistically using Wald’s type II tests (*X*^2^ estimates that have a *P*<0.05 are boldfaced). The table also includes (B) estimates of the crop and country random effects with their respective 95% confidence intervals, (C) estimates of the percentage of variation in yield decline explained by the fixed factors included in each model (*i.e.*, marginal *R*^2^) and by each model as a whole (*i.e.*, conditional *R*^2^), and (D) comparisons with homologous phylogenetic GLMM models and estimates of phylogenetic signal in models’ residuals.

**(A) Fixed effects**
|  | GLMM_0 |  |  | GLMM_1a |  | GLMM_1b |  | GLMM_2 |  | GLMM_3 |  |
| --- | --- | --- | --- | --- | --- | --- | --- | --- | --- | --- | --- |
| | Df | $\chi^2$ | *P* | $\chi^2$ | *P* | $\chi^2$ | *P* | $\chi^2$ | *P* | $\chi^2$ | *P* |
| Pollinator dependence (P) | 2 | – | – | **6.71** | 0.035 | – | – | 2.54 | 0.28 | 2.08 | 0.35 |
| Growth form (G) | 2 | – | – | – | – | **15.95** | 0.00034 | **11.64** | 0.0029 | **11.93** | 0.0026 |
| Region (R) | 4 | – | – | **26.37** | $<10^{-4}$ | **26.16** | $<10^{-4}$ | **26.39** | $<10^{-4}$ | **24.75** | $<10^{-4}$ |
| P x G | 4 | – | – | – | – | – | – | – | – | 4.06 | 0.40 |
| P x R | 8 | – | – | – | – | – | – | – | – | **19.53** | 0.012 |
| G x R | 8 | – | – | – | – | – | – | – | – | **28.53** | 0.00038 |
| Area_crop | 1 | – | – | **37.51** | $<10^{-8}$ | **36.54** | $<10^{-8}$ | **34.26** | $<10^{-8}$ | **35.66** | $<10^{-8}$ |
| Area_country | 1 | – | – | **26.19** | $<10^{-7}$ | **25.55** | $<10^{-6}$ | **25.76** | $<10^{-6}$ | **26.40** | $<10^{-6}$ |
| Estimate |  | Slope | SE | Slope | SE | Slope | SE | Slope | SE | Slope | SE |
| Area_crop |  | – | – | -0.481 | 0.079 | -0.461 | 0.076 | -0.445 | 0.076 | -0.467 | 0.078 |
| Area_country |  | – | – | -0.359 | 0.070 | -0.353 | 0.070 | -0.354 | 0.070 | -0.362 | 0.071 |

**(B) Random effects**
|  | GLMM_0 |  | GLMM_1a |  | GLMM_1b |  | GLMM_2 |  | GLMM_3 |  |
| --- | --- | --- | --- | --- | --- | --- | --- | --- | --- | --- |
|  | Variance | 95%CI | Variance | 95%CI | Variance | 95%CI | Variance | 95%CI | Variance | 95%CI |
| Crop | 0.466 | (0.30-0.71) | 0.248 | (0.15-0.41) | 0.217 | (0.13-0.36) | 0.204 | (0.12-0.35) | 0.197 | (0.12-0.33) |
| Country | 0.695 | (0.49-0.99) | 0.455 | (0.31-0.66) | 0.448 | (0.31-0.66) | 0.449 | (0.31-0.66) | 0.453 | (0.31-0.67) |

**(C) Model explanatory power**
|  | GLMM_0 | GLMM_1a | GLMM_1b | GLMM_2 | GLMM_3 |
| --- | --- | --- | --- | --- | --- |
| Marginal $R^2$ | 0.000 | 0.101 | 0.110 | 0.111 | 0.138 |
| Conditional $R^2$ | 0.261 | 0.259 | 0.260 | 0.258 | 0.280 |

**(D) Phylogenetic effects**
|  | GLMM_0 |  | GLMM_1a |  | GLMM_1b |  | GLMM_2 |  | GLMM_3 |  |
| --- | --- | --- | --- | --- | --- | --- | --- | --- | --- | --- |
| | AIC | ΔAIC | AIC | ΔAIC | AIC | ΔAIC | AIC | ΔAIC | AIC | ΔAIC |
| GLMM | 4229.11 |  | 4148.85 |  | 4140.31 |  | 4141.82 |  | 4130.16 |  |
| Phylo-GLMM | 4232.34 | 3.23 | 4152.43 | 3.58 | 4147.19 | 6.88 | 4150.26 | 8.44 | 4138.74 | 8.58 |
| Phylogenetic signal in the model residuals | Estimate | $P$ | Estimate | $P$ | Estimate | $P$ | Estimate | $P$ | Estimate | $P$ |
| $K$ | $4.51 \cdot 10^{-7}$ | 0.35 | $6.79 \cdot 10^{-7}$ | 0.13 | $6.39 \cdot 10^{-7}$ | 0.15 | $6.28 \cdot 10^{-7}$ | 0.15 | $6.28 \cdot 10^{-7}$ | 0.15 |

### The importance of pollinator dependence and crop growth form

Pollinator dependence and plant growth form were related traits. Pollinator dependence and growth form were associated across the whole set of 136 crops (*X*^2^=26.3, df=4, *P*<0.001). Specifically, the yield of about 64.55% of all herbaceous crops did not depend on pollinators, whereas only 12.66% of crops with that growth form depended highly on pollinators. On the other hand, only 24.39% of all tree crops were pollinator independent, whereas 48.78% were highly dependent on pollinators (Fig. 3). Shrub crops showed intermediate values, with 37.5 and 18.75% of these crops exhibiting no or high pollinator dependence, respectively. Considering crops cultivated exclusively for their reproductive organs (105 crops) did not alter this association (*X*^2^=18.4, df=4, *P*=0.001), with 18.18, 25.0, and 52.62% of herb, shrub, and tree crops depending highly on pollinators, respectively.

**Fig. 3.**
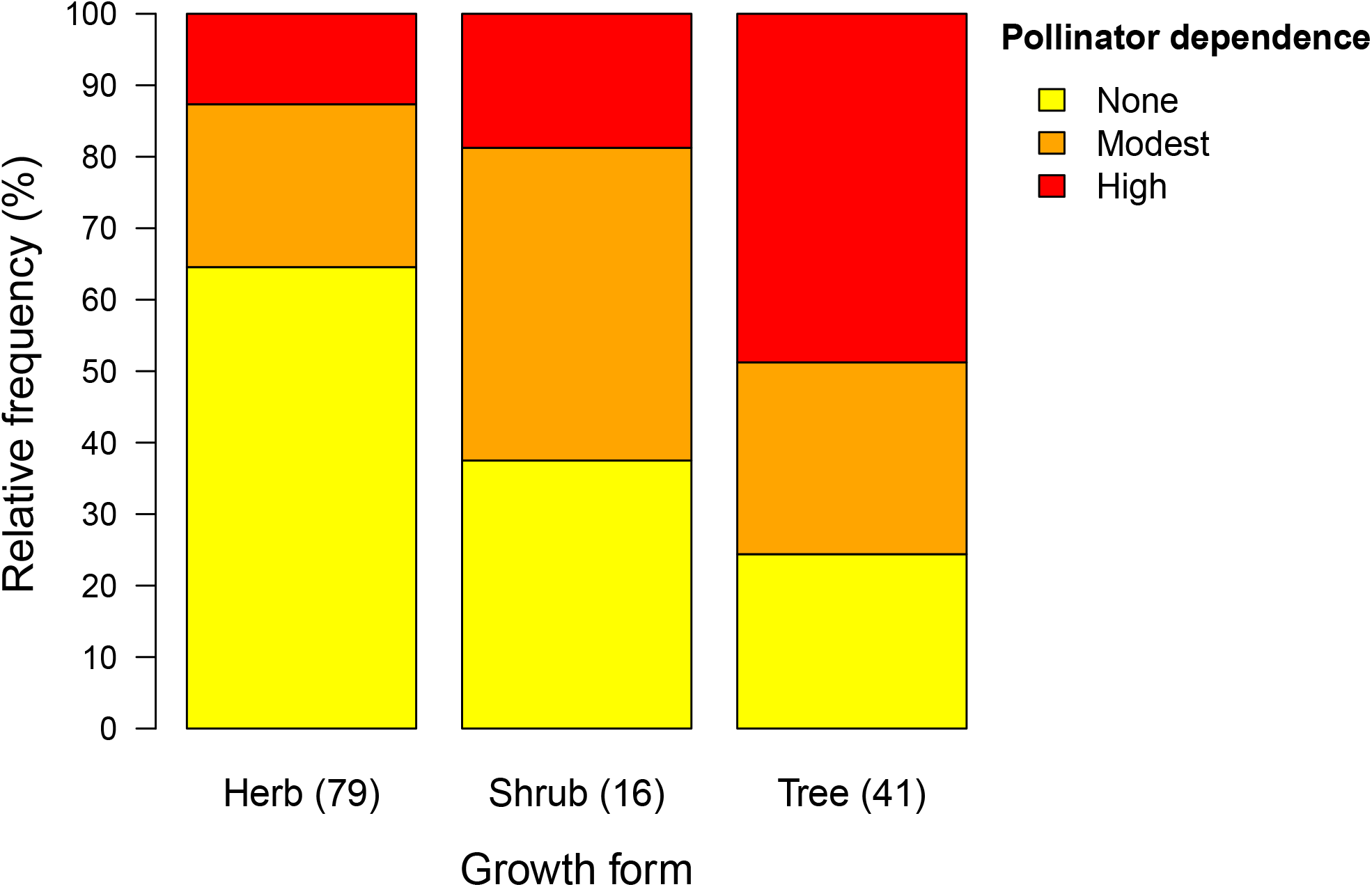
Relative frequency of crops with none, modest, and high dependence on pollinators across growth-form categories (*i.e.*, herb, shrub, and tree crops). In parentheses, the number of crops in each growth-form category.

High pollinator dependence and the tree growth form were both plant characteristics associated with an enhanced probability of yield decline when considered separately (models GLMM_1a and GLMM_1b; Fig. 4 and Table 1). Particularly, these two traits increased the probability of yield decline by ∼40 and ∼60%, in comparison with pollinator-independent and herbaceous crops, respectively (Fig. 4). In the absence of the other predictive focal factor, pollinator dependence and growth form accounted for 5.6 and 12.9% of the among-crop variance in yield decline, respectively (Fig. 5). However, the effect of pollinator dependence almost vanished, whereas the effect of plant growth form persisted, when both factors were included in the same model (GLMM_2; Figs. 4-5 and Table 1). These statistical results were robust to different data manipulations, such as excluding the modal years 1961 and 2020 (Fig. 1) in the estimation of yield decline or, as in Deguines et al. (2014), considering pollinator dependence as a numerical rather than as a categorical variable (Table S3).

**Fig. 4.**
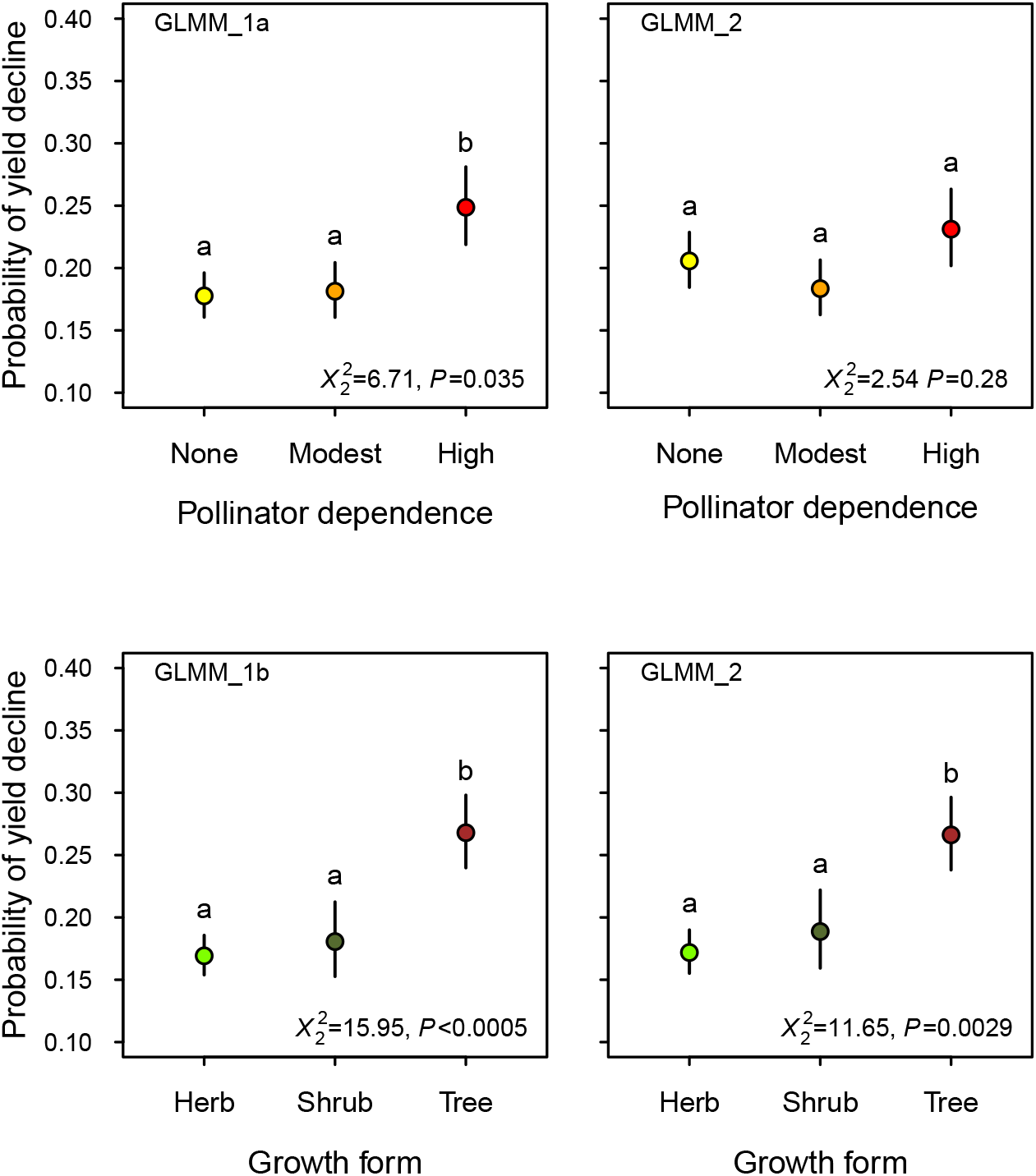
Probability of yield decline in relation to pollinator dependence and growth form. The figure depicts the mean estimates (+/- 1 SE) of the proportion of crops in each pollinator dependence and growth-form category with negative average growth rates during the period 1961-2020. The left two panels (GLMM_1a and GLMM_1b) show the results of the effect of each factor when the other factor was not included in the model (*i.e.*, model GLMM_1a tested the effect of pollinator dependence and model GLMM_1b tested the effect of growth form), whereas the right two panels the results of each factor after accounting for the confounding effect of the other factor (model GLMM_2). Means with the same letter do not provide evidence of statistical differences at the level of α = 0.05 according to a pairwise Tukey’s a posteriori test. Wald’s type II test results are provided (see also Table 1).

**Fig. 5.**
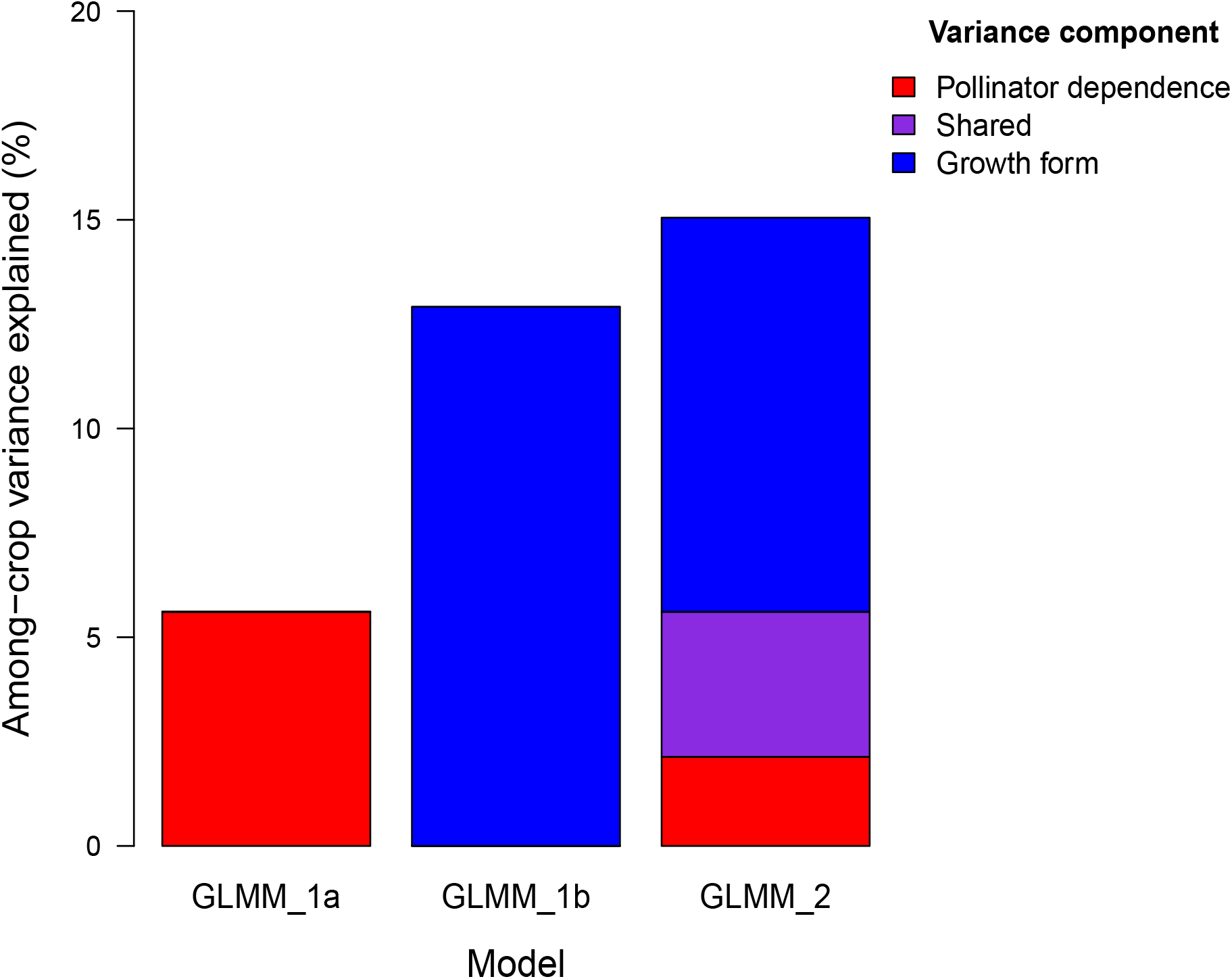
Percentage of among-crop variance in yield decline explained by pollinator dependence and growth form in the absence of the other focal factor as estimated from models GLMM_1a and GLMM_1b, respectively, and the independent and shared percentage of the among-crop variance explained by these two focal factors as estimated from model GLMM_2. All these components of variance exclude any variation that could also be accounted by differences among crops in cultivated area (Table 1).

We found no evidence of an interaction between pollinator dependence and growth form, but there was evidence of regional differences in the effect of pollinator dependence on yield decline (GLMM_3; Table 1). This interaction effect was mostly attributed to Asia where, contrary to expectation, the estimated probability of yield decline was somewhat lower among pollinator-dependent crops (Fig. 6). There was also an indication of regional differences in the effect of growth form on the probability of yield decline (GLMM_3; Table 1). Results show that a decline in the yield of tree crops was strongest in Europe and Oceania followed by Asia and the Americas. Africa was the only region where yield decline was not clearly associated with tree crops (Fig. 7).

**Fig. 6.**
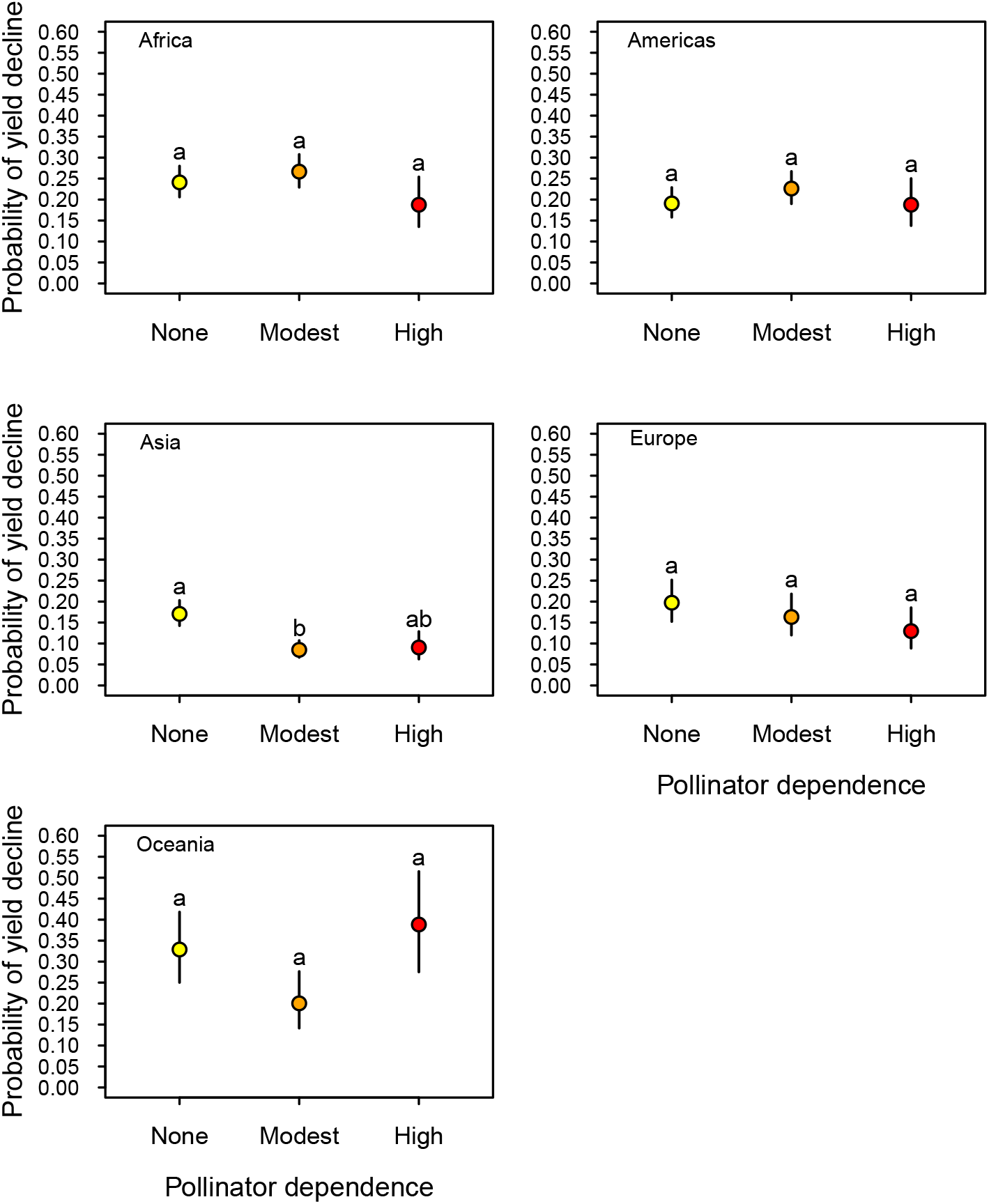
Probability of yield decline in relation to pollinator dependence by geographical region. The figure depicts the mean estimates (+/- 1 SE) of the proportion of crops in each pollinator-dependence category with negative average growth rates in each region during the period 1961­2020 according to model GLMM_3. Means with the same letter in each panel do not provide evidence of statistical differences at the level of α = 0.05 according to a pairwise Tukey’s a posteriori test.

**Fig. 7.**
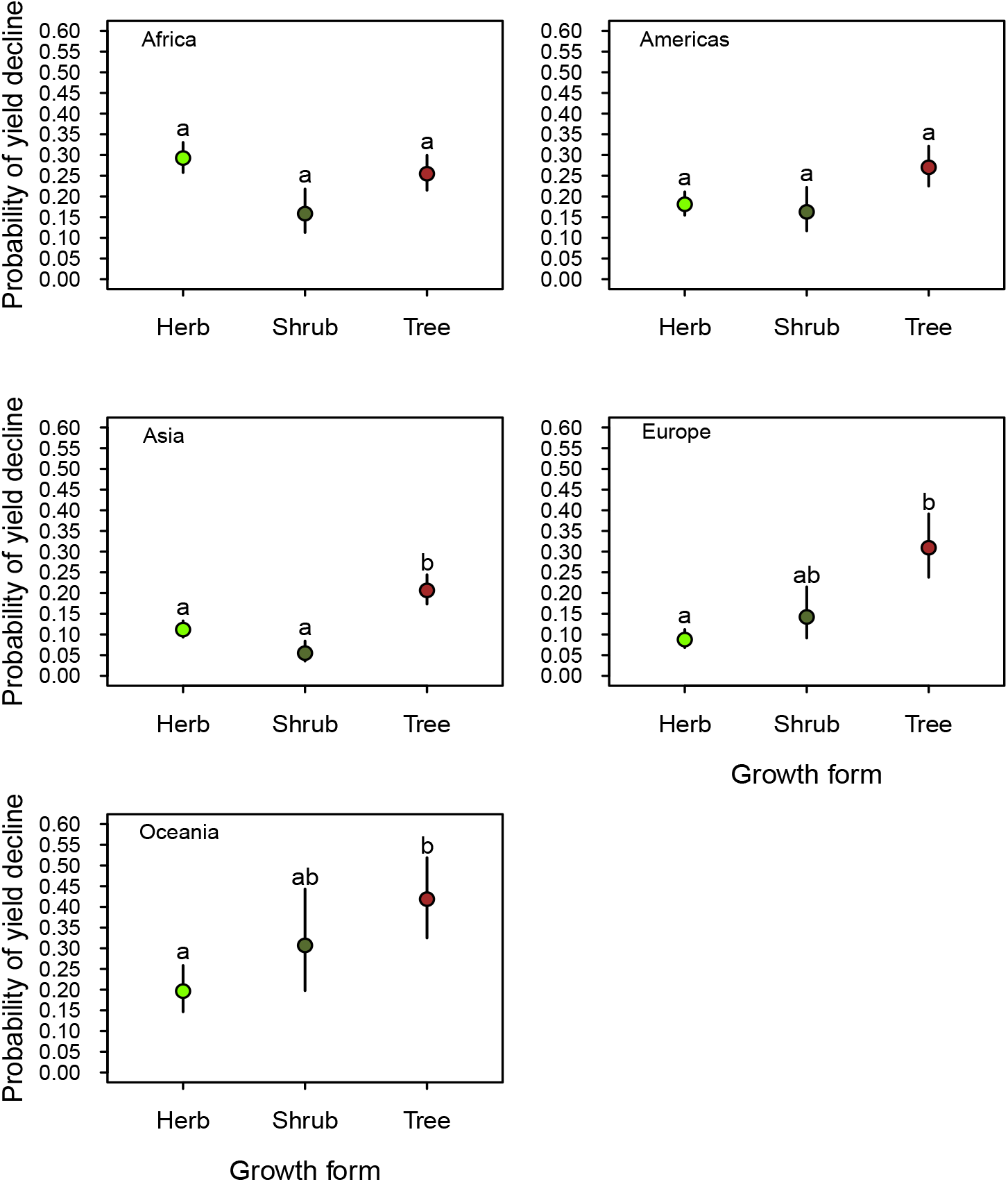
Probability of yield decline in relation to crop growth form by geographical region. The figure depicts the mean estimates (+/- 1 SE) of the proportion of crops in each growth-form category with negative average growth rates in each region during the period 1961-2020 according to model GLMM_3. Means with the same letter in each panel do not provide evidence of statistical differences at the level of α = 0.05 according to a pairwise Tukey’s a posteriori test.

### Accounting for potentially confounding factors

We found a non-linear strong decrease in the probability of yield decline with increasing total cultivation area per crop and country (Table 1, Fig. S5). However, the relations between the focal variables, pollinator dependence and plant growth form, and the probability of decline reported were independent of these effects, as these area variables were included as fixed-factor effects in our models. Also, data analyses revealed no evidence for differences in the estimated probability of yield decline between crops cultivated for their reproductive *vs*. vegetative parts (least-square means [-1SE/+1SE]=0.18 [-0.023/+0.026] *vs*. 0.14 [-0.021/+0.025]; *X*^2^=1.89, *P*=0.17). Despite the association between pollinator dependence and plant growth form, there was no evidence that multicollinearity could introduce biases in the assessment of any of the factors and covariates included in our analyses (all adjusted GVIFs <1.1). Last, non-phylogenetic models provided a better fit to the data than phylogenetically-explicit models based on AIC criteria (ΔAICs> 3), whereas there was no evidence of phylogenetic structure in the crop-averaged residuals as revealed by the low magnitude of Blomberg’s *K* signal (Table 1).

## DISCUSSION

Crop yield declines are widespread, with approximately one-quarter of all crop × country yield trends in our dataset exhibiting signs of decline. However, the probability of yield decline was highly heterogeneous among crops and countries. Part of that heterogeneity was explained by cropś dependence on pollinators when this factor was considered in isolation. But this association mostly disappeared when accounting for the stronger effect of plant growth form. Therefore, given the association between high pollinator dependence and the tree growth form in crops, the effect of pollinator dependence on yield declines seems to be a side effect of a more direct association between the probability of yield decline and plant growth form. In particular, we reveal that tree crops are more likely to experience yield declines.

The association between growth form and pollinator dependence across crops we report here (Fig. 3) has relevant conceptual and practical implications. Despite thousands of years of plant domestication, patterns of covariation between vegetative and reproductive traits in crop plants still reflect the contrasting life-history strategies that characterize wild plants at large. Both wild flowering plants and crops show associations between growth form, lifespan, flower size and numbers, outcrossing, and, as a consequence, the extent of pollinator dependence. Trees, in particular, are long-lived, produce lots of relatively small flowers, and show high frequencies of self-incompatibility that enforce outcrossing, which results in high pollinator dependence (Friedman 2020; Cunha and Aizen 2023; Lanuza et al. 2023). Because somatic mutations may be passed on to seeds due to a lack of a segregated germline in plants, mutational load accumulation with increasing perenniality is probably the ultimate driver behind these associations, (Klekowski 1988; Schoen and Schultz 2019). Thus, selection for complete pollinator independence in pollinator-dependent tree crops may be unfeasible (Sáez et al. 2020). More generally, the above patterns of covariation may limit the phenotypic space that can be explored via either artificial selection or genetic engineering (Milla et al. 2018; Garibaldi et al. 2021; Cunha et al. 2023).

Pollinators are declining worldwide as a part of the ongoing global biodiversity crisis (Potts et al. 2010; Zattara and Aizen 2021). This trend threatens not only the yield of hundreds of crops but also the reproduction of thousands of wild plant species (Rodger et al. 2021). In particular, diverse pollinator assemblages play a crucial role in maintaining high yields of many common, nutritionally important, and economically valuable crops, such as coffee, stone-fruit crops, and cucurbits (Garibaldi et al. 2013). Although dozens of studies have shown that spatial and temporal local declines in wild pollinator populations negatively affect the yield of many pollinator-dependent populations, evidence that pollinator decline has impacted crop yields globally has remained elusive (Aizen et al. 2008, 2022). For instance, global data indicate that yield growth rates and their stability seem to decrease with increasing pollinator dependence (Garibaldi et al. 2011a). However, these findings cannot be taken as unequivocal evidence of the impact of pollinator decline despite widespread pollination limitation (Ashman et al. 2004), as they may be more strongly and proximately influenced by mate than pollinator availability (Harder and Aizen 2010; Sáez et al. 2022) or by other correlated factors as shown here. In addition, previous analyses of global data have failed to find a deceleration in yield growth with increasing pollinator dependence (Aizen et al. 2022). In our study, we did not find evidence to support the proposal that a crop’s pollinator dependence is a proximal driver of yield decline after accounting for the association between growth form and pollinator dependence. Asia was the only region where there was evidence that the probability of yield decline changes with pollinator dependence irrespective of growth form. However, the observed pattern was contrary to expectations, suggesting that pollination management, including human hand pollination (Wurz et al. 2021), might have counteracted potential impacts of pollinator decline in that region. Therefore, even though evidence implies that pollinator decline is occurring at small as well as continental spatial scales, this phenomenon does not seem to have affected crop yield globally. This lack of evidence does not rule out the possibility that pollinator decline may be affecting the yield of particular crops in some areas, but it implies that pollinator dependence cannot be considered the primary driver of yield decline in most pollinator-dependent crops. Breeding of new less pollinator-dependent varieties of typically highly pollinator-dependent crops, such as almonds (Sáez et al. 2020), and more efficient management of crop pollination (Mueller et al. 2012; Röös et al. 2018) might be offsetting the effect of increasing pollination deficits due to dwindling pollinator populations.

Beyond logging and habitat destruction, tree mortality rates have increased in forests around the world over the past few decades, likely as a result of climate change and associated stressors, such as higher occurrences of insect outbreaks, wildfires, heat waves, and frosts (Allen et al. 2010; BGCI 2021). Sustained declines in fruit and seed production, the most common harvest of tree crops, precede tree death (Pesendorfer et al. 2019). Thus, decreasing tree crop yields over time could signal the impacts of climate change. In addition, interannual yield variation is higher in woody than in herbaceous crops, independent of pollinator dependence and harvest organ (Gleiser et al. 2021). Although comparative studies on the susceptibility of different plant growth forms to climate change are lacking, a recent study has revealed that herbaceous plants have been more tolerant to frost than woody plants over evolutionary time (Klimeš et al. 2022). This differential susceptibility could be due to frost-caused xylem embolism and cavitation, which could trigger wood dieback (Martínez-Vilalta and Pockman 2002; Mayr et al. 2003). Because one of the consequences of climate change is an increasing incidence of spring-frost damage due to advances in spring phenology (Lamichhane 2021), this could be one of the contributing factors behind the differential yield decline of tree crops compared to that of herbaceous crops. Even though frosts can ruin the harvest of an annual herbaceous crop, re-sowing during the same or the year following the crop’s failure would restore prior yields. Also, the short lifespan and high relevance of several herbaceous plants as basic staple crops make their genetic manipulation more short-term amenable and profitable than most long-lived perennial plants (McCown 2000), resulting in the production of new crop varieties that can adapt rapidly to a changing climate (Henry 2020). Interestingly, the two crops showing the lowest probability of yield decline were maize and wheat (Fig. S2), two of the most genetically engineered crops (Takeda and Matsuoka 2008).

Our findings suggest that the association between pollinator dependence and the probability of yield decline is largely explained by its partial association with growth form. Specifically, we found a higher frequency of high pollinator dependence among tree crops, confounding the effects of pollinator dependence and growth form. However, this association was not as complete as to prevent evaluation of the independent effect of each of these two factors on the probability of yield decline. In addition to showing that the relationship between growth form and the probability of yield decline is relatively strong, we found that the increase in the probability of yield decline among tree crops was consistent across continents, except in Africa. Africa was the continent with the highest probability of crop yield declines, which might relate to a combination of events, including an increasing occurrence of dry spells, and political, social, and economic factors that affect the agricultural sector as a whole (*e.g.*, absence of significant irrigation infrastructure), irrespective of plant growth form (Ray et al. 2012). The total cultivated area per country and crop was also a potential confounding factor. The most widely cultivated crops were both herbaceous and pollinator independent, which showed the lowest probabilities of yield declines. However, the reported association between plant growth form and the probability of yield decline accounts for this potential confounding effect. In addition, the fact that most tree crops are cultivated for either their seeds or fruits may also confound the effect of the type of organ harvested (vegetative *vs.* reproductive) with the effect of plant growth form. Yet, we did not find evidence that the type of organ harvested influences the probability of decline to any significant extent. Finally, we can discard any effects of unmeasured phylogenetically-conserved factors on the probability of yield decline, given the worse goodness of fit of the phylogenetically-explicit models and the lack of phylogenetic signal in model residuals. Therefore, the growth form of a crop plant seems to connect more proximately with the likelihood of exhibiting a long-term decline in yield than any other of the factors studied.

Although we are interpreting the association between growth form and the probability of yield decline in the context of climate change, it is essential to explore other explanations for this relationship that are independent of the environmental context. One possible explanation is that potentially decreasing market prices may have discouraged the cultivation of tree crops and the proper management of existing cultivated fields. However, the area cultivated with fruit and seed crops has been increasing steadily for decades (Aizen et al. 2022), with market prices for those crops that are several times higher than those of cereals and most other crops cultivated for their vegetative parts (Gallai et al. 2009). Another potential explanation could be the replacement of slow-growing tree crops by fast-growing and fast-cash herbaceous crops, which may have left remaining fields of at least some tree crops unattended, or restricted those crops to less productive marginal areas. For example, the rapid expansion of soybean cultivation in several countries of the Americas in the last decades has impacted the diversity of cultivated crops. However, this replacement cannot be considered a global phenomenon, and in some regions such as Europe and countries elsewhere, the replacement seems to have been in the opposite direction (Aizen et al. 2019). In conclusion, the proximate factors explaining the relationship between growth form and yield declines need to be investigated more in depth. However, climate change seems to be a plausible overarching phenomenon behind the reported association.

## CONCLUDING REMARKS

The deceleration in yield growth is a complex phenomenon with multiple causes, including limitations in crop improvement and diminishing yield returns to increasing external subsidies such as irrigation, fertilizers, and pesticides (Ray et al. 2012). However, yield decline, rather than just yield growth deceleration, is also likely to reflect the consequences of widespread environmental degradation and not just the reach of human management skills. This is particularly so when negative growth rates are also related to biological crop traits like growth form, as reported here. Our study also found that yield decline is widespread but exhibits high geographic variability. While further research is needed to understand why yield decline is more severe in some countries and regions, we found that plant growth form, rather than a crop’s pollinator dependence, is a more proximate factor explaining variation in yield decline at regional scales. This highlights the importance of not considering any single factor in isolation but contrasting the explanatory power of each factor with other correlated factors in observational studies. For example, we might have reached a misleading conclusion about the relationship between pollinator dependence and yield decline if we did not consider that pollinator dependence is associated with growth form. In particular, our study revealed a differential incidence of yield decline among tree crops compared to crops with other growth forms, paralleling reports of widespread mass tree mortality associated with climate change. While climate change can provide a general explanation for this association, more research is needed to understand the physiological mechanisms behind it for proper crop management and breeding. Beyond highlighting the need for this type of follow-up research, the reported association between plant growth form and yield decline adds to the evidence of the potential hazards of climate change for food security.

## Acknowledgments

The authors acknowledge C.L. Morales, L.D. Harder, and A. Sáez for helpful discussions, and I. Bartomeus, N. Deguines, and an anonymous reviewer for useful comments and suggestions on a previous version of this contribution.

## Authors contribution

M.A.A, G.G., T.K, and R.M. conceived the study. M.A.A. and G.G. compiled and analyzed the data. R.M. provided feedback on the analyses. M.A.A. wrote the paper with input from all authors.

## Data and script accessibility

Data and scripts for this publication are available on the Zenodo Repository (Aizen et al. 2023): https://doi.org/10.5281/zenodo.8206104.

## Appendix

**Fig. S1.**
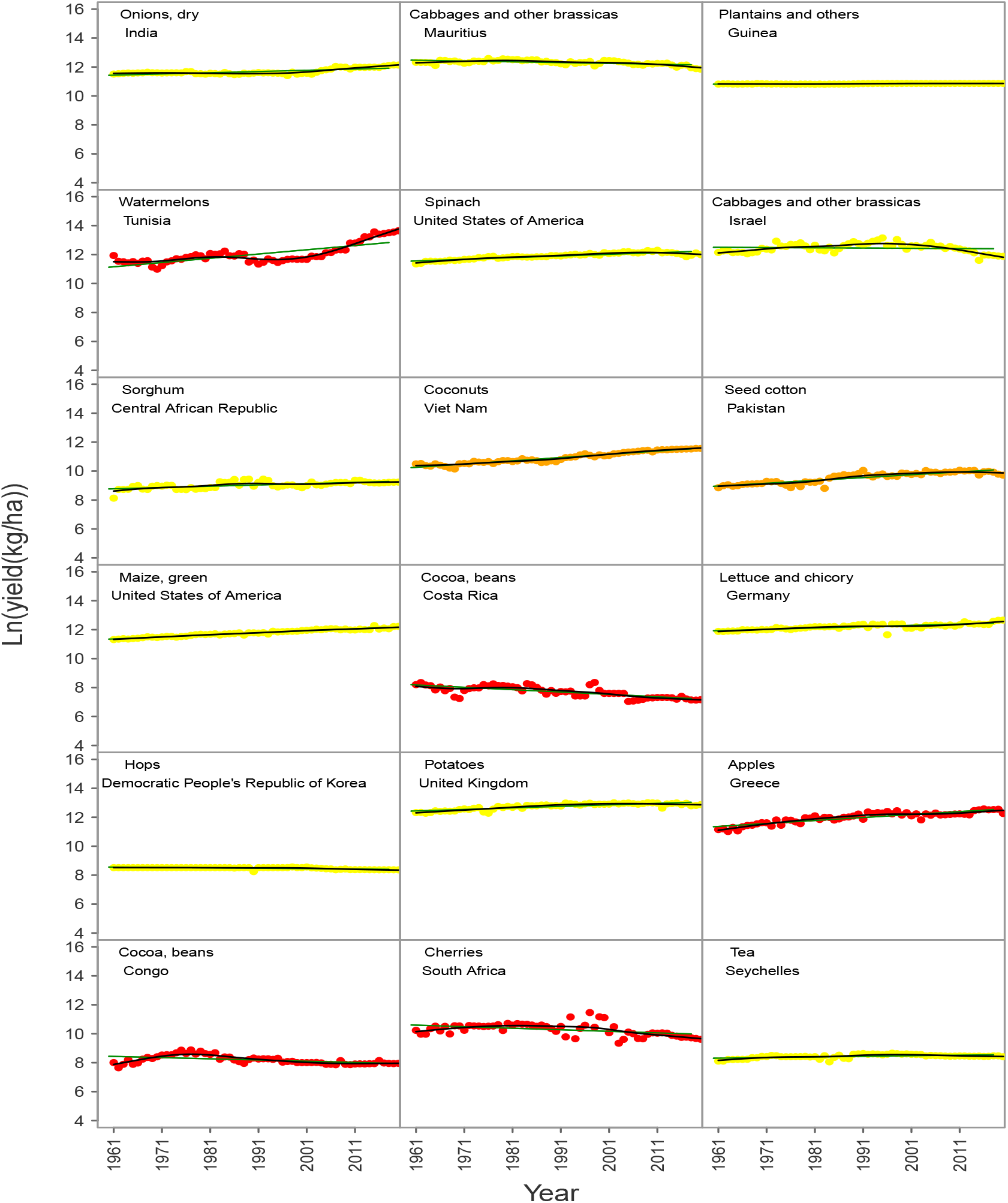
A randomly-chosen sample of yield trends (1961-2020) for different crops in different countties. Point colors represent different categories of pollinator dependence (i.e., none=yellow, modest=orange, high=red). The difference in the natural logarithm of yield between any two given years estimates the relative yield growth during that period, whereas the slope of the regression line of (natural-log) yield as a function of year estimates the annual growth rate in yield. The sign of the regression was used to characterize whether the yield of a given crop in a given country showed evidence of long-term decline or not. For instance, maize in the USA showed no evidence of decline, whereas cocoa in Costa Rica did. Probability of yield decline

**Fig. S2.**
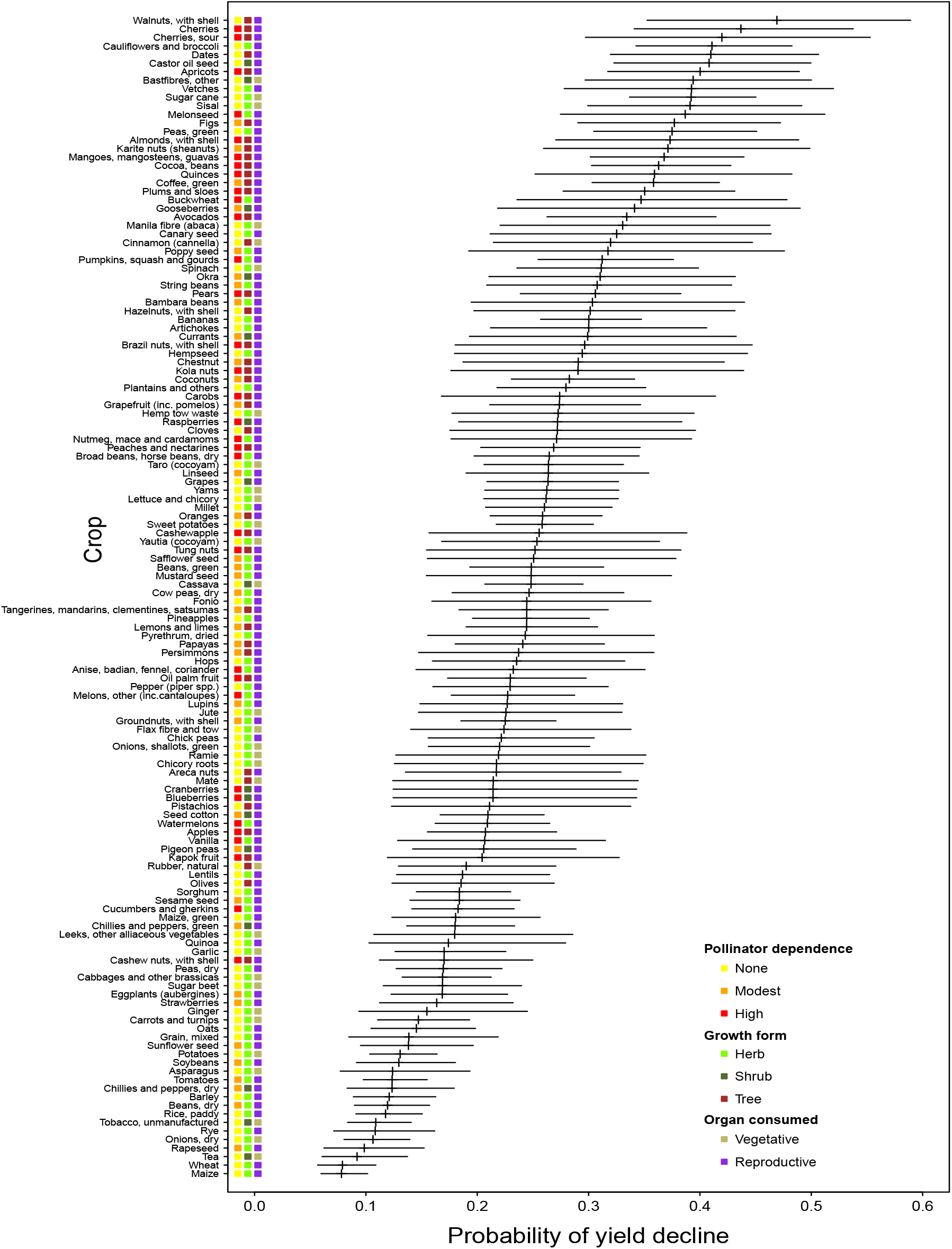
Variation in the probability of yield decline among crops. As a crop estimate of the probability of yield decline, the figure depicts the expected proportion of countries (+/- 1SE) in which each crop showed average growth rates in yield <0 over the period 1961-2020. Estimates are based on model GLMM_0 (Table 1). Crops are classified based on their pollinator dependence, growth form, and type of organ consumed.

**Fig. S3.**
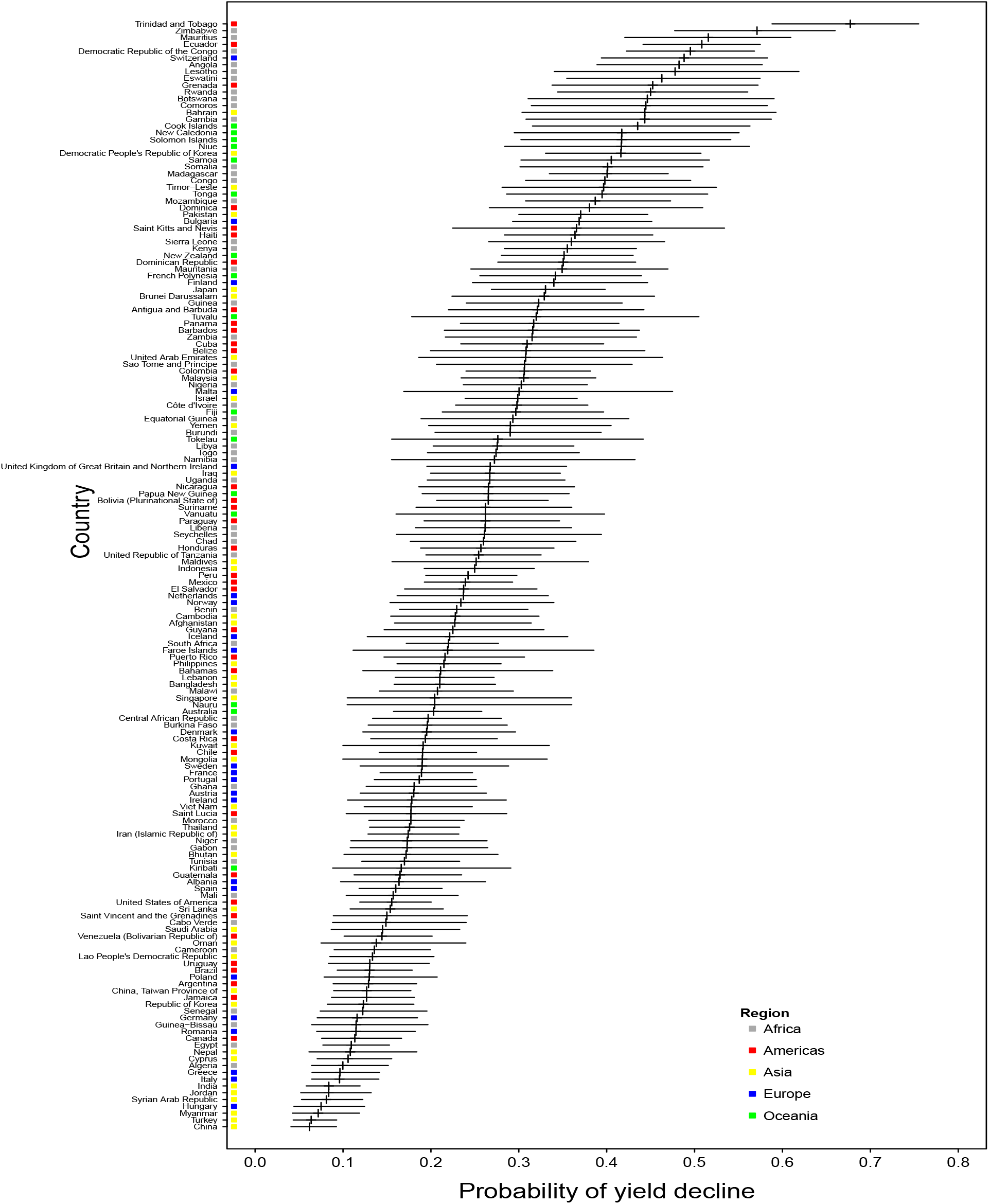
Variation in the probability of yield decline among countries. As a country estimate of the probability of yield decline, the figure depicts the estimated proportion of crops (+/- 1SE) showing average growth rates in yield <0 for each country over the period 1961-2020. Estimates are based on model GLMM_0 (Table 1). Countries are classified according to geographical region.

**Fig. S4.**
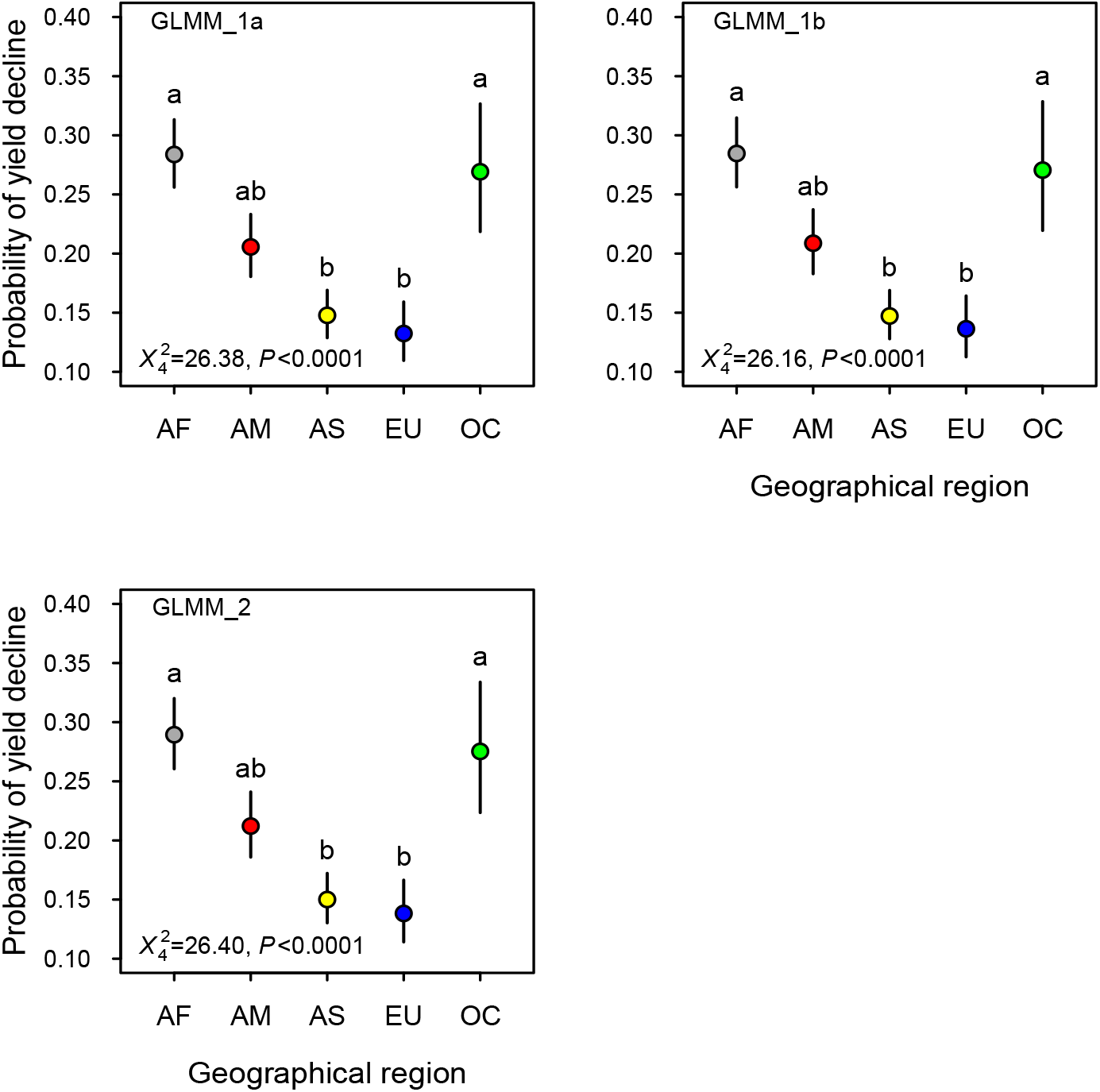
Probability of yield decline by geographical region as estimated by the main effects models. The figure depicts the mean estimates (+/-1 SE) of the proportion of crops showing negative average growth rates for countties in each region of the world (i.e., AF, Africa; AM, the Americas; AS, Asia; EU, Europe; OC, Oceania) during the period 1961-2020 according to the main-effects GLMMs that included this factor (i.e., GLMM_la, GLMM_lb, GLMM_2). Means with the same letter do not provide evidence of statistical differences at the level of α = 0.05 according to a pairwise Tukey’s a posteriori test. Wald’s type II test results are provided (see also Table 1).

**Fig. S5.**
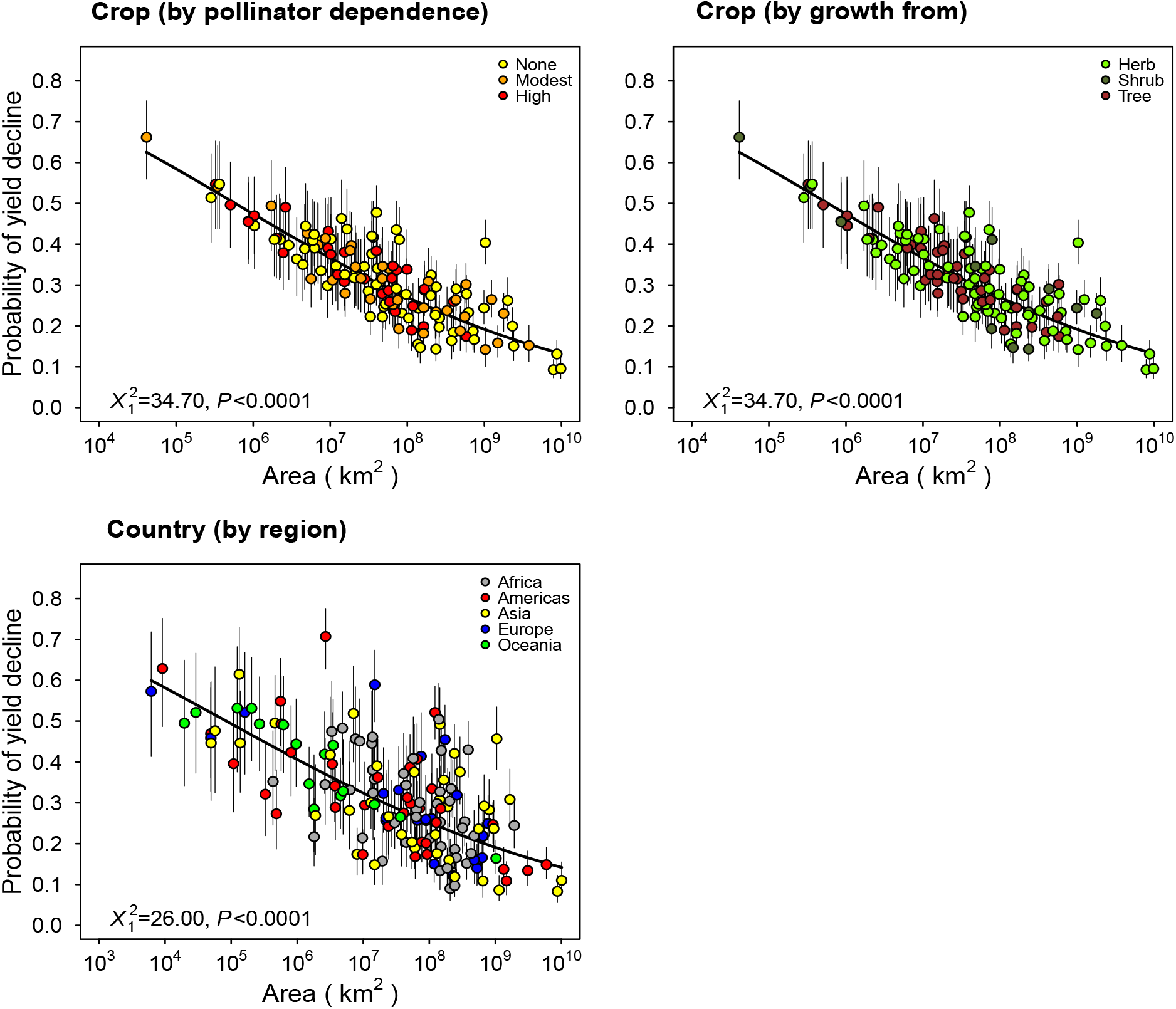
Probability of yield decline and cumulative cultivated area per crop and country. The figure depicts the mean estimates (+/-1 SE) of the proportion of countties in which a given crop showed negative average growth rates (upper two panels) or the proportion of crops in a given country showing negative average growth rates (lower panel) as a function of the cumulative cultivated area (1961-2020) per crop or country, respectively. Estimates were exttacted from model GLMM_2, even though the effect of area was very similar in all models that included the effects of cultivated area per crop and country (Table 1). In the upper panels, crops were categorized by pollinator dependence (left) and growth form (right), and in the lower panel countties were categorized by geographical region.

**Table S1.**
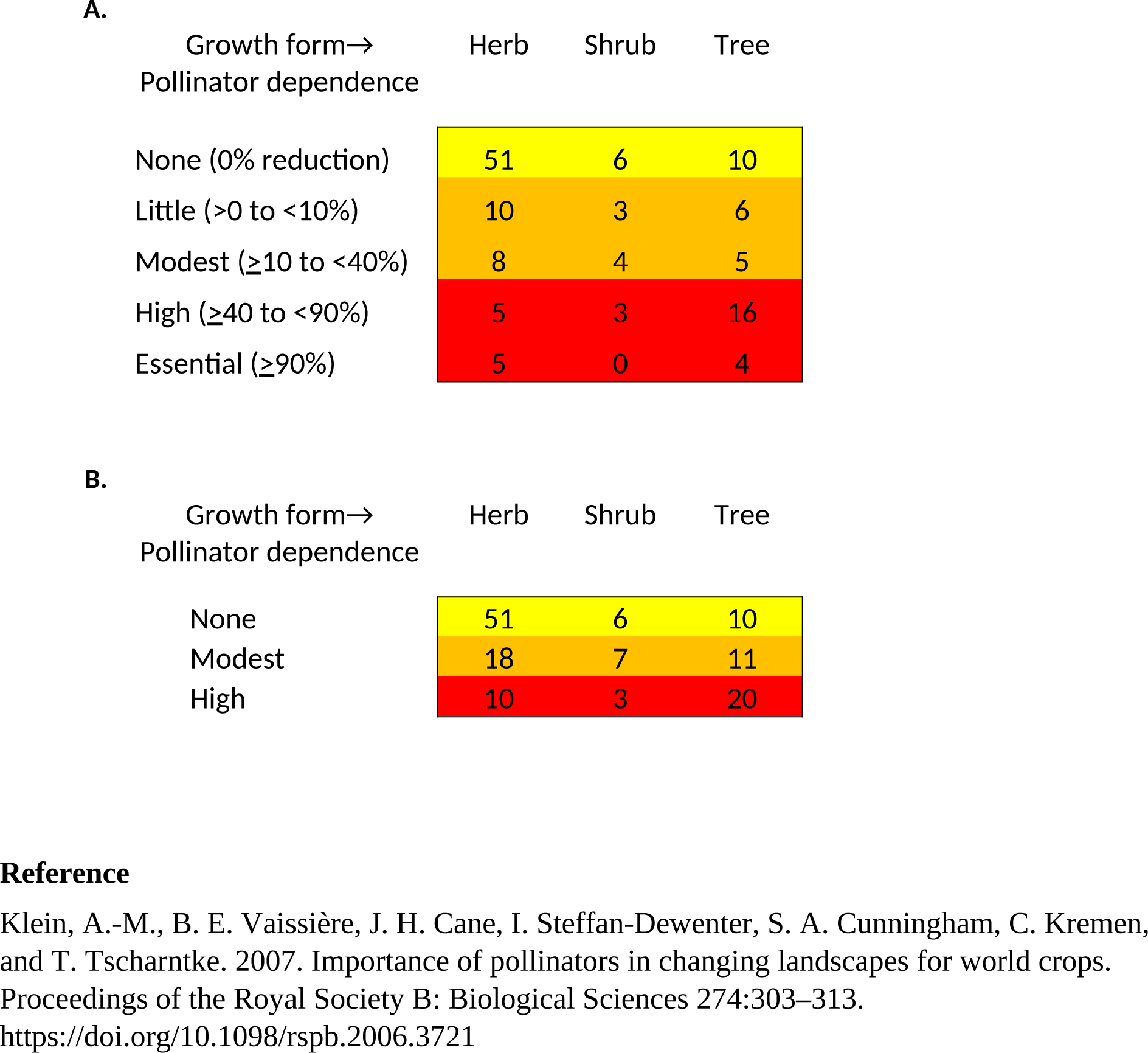
Two-way frequency tables showing the number of crops in each combined category of pollinator dependence x growth form according to (A) the five pollinator-dependence levels defined by Klein et al. (2007), and (B) the three pollinator-dependence levels considered in this contribution resulting from the lumping of the levels highlighted with the same color as in (A).

**Table S2.**
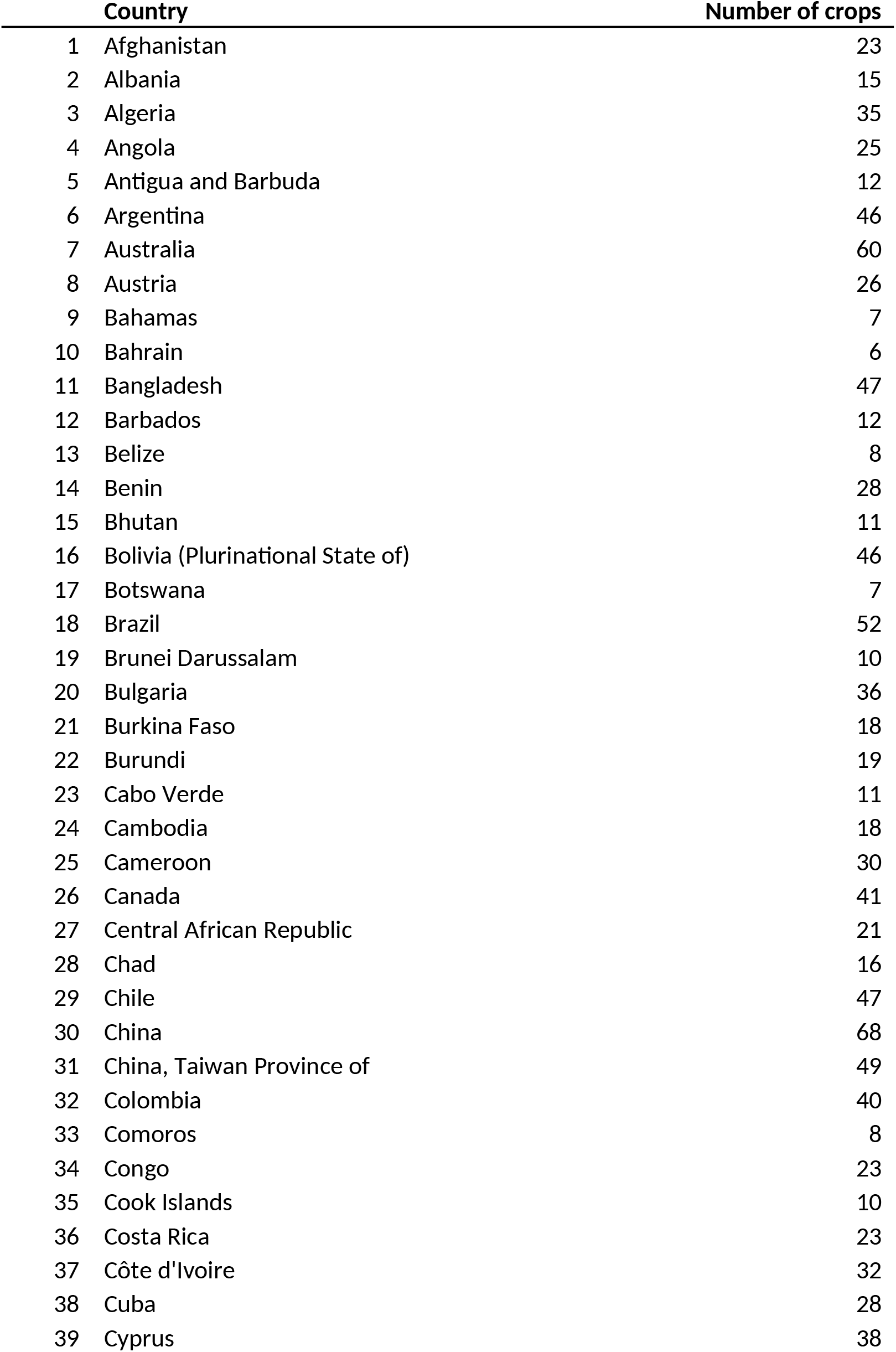
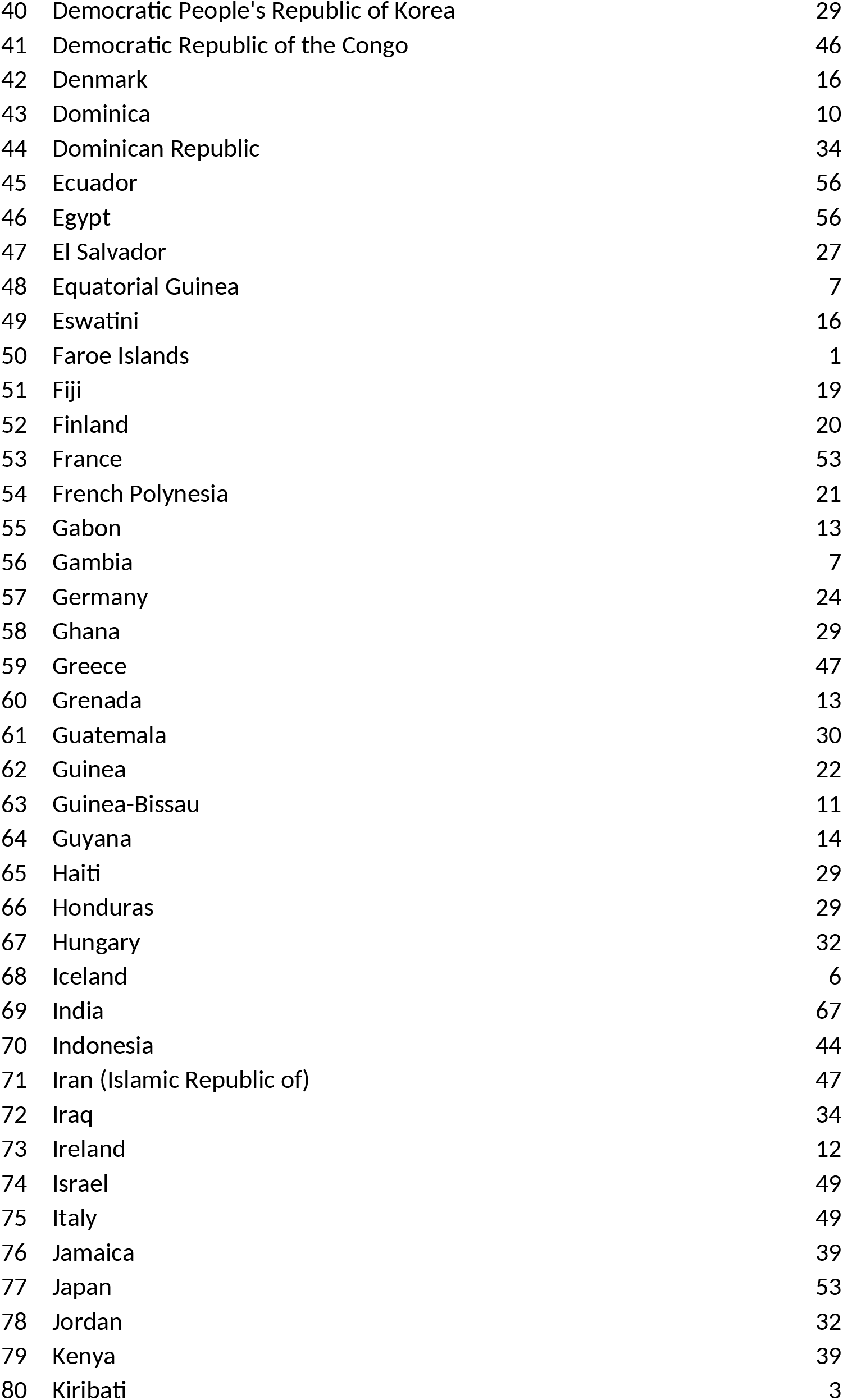
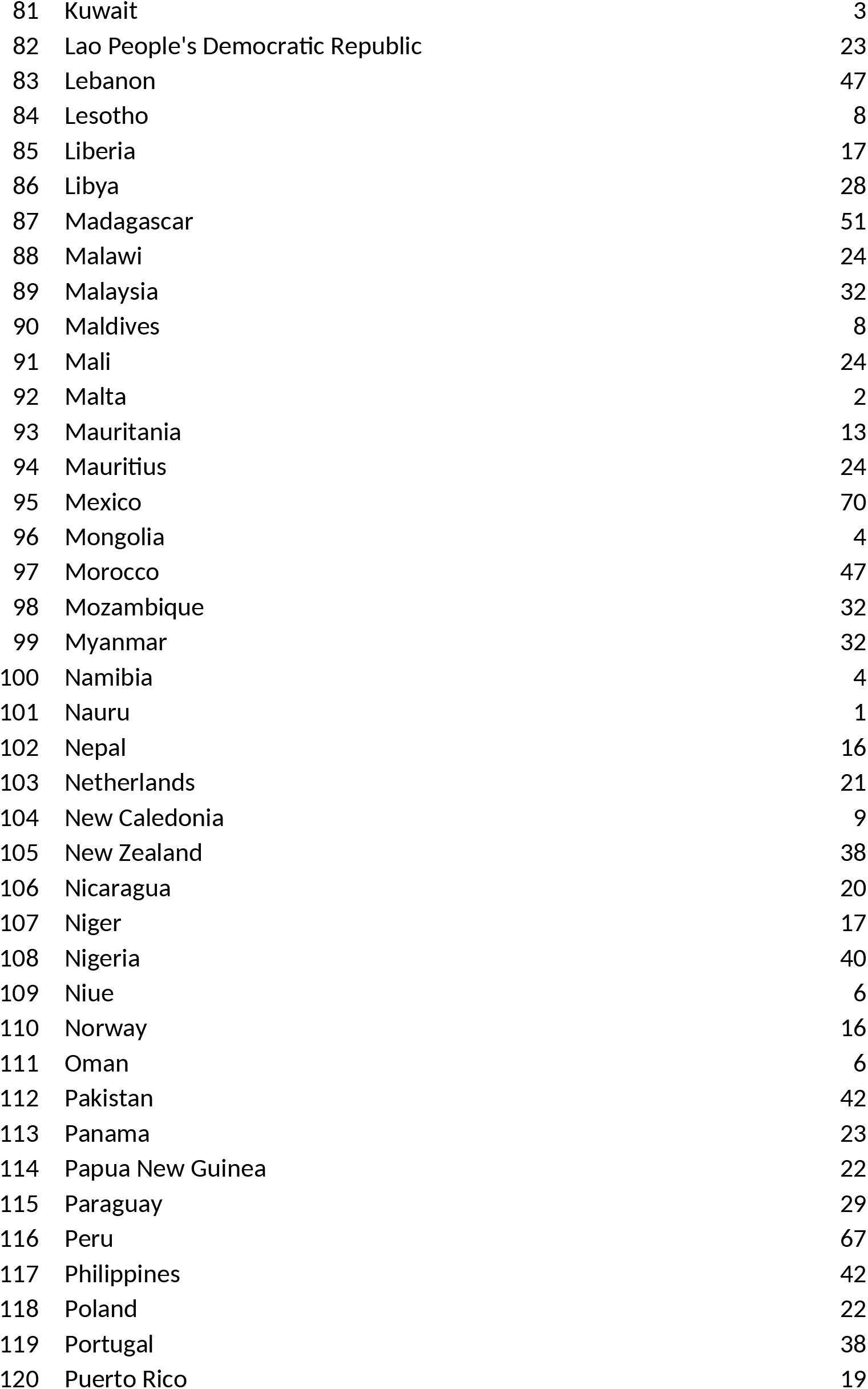
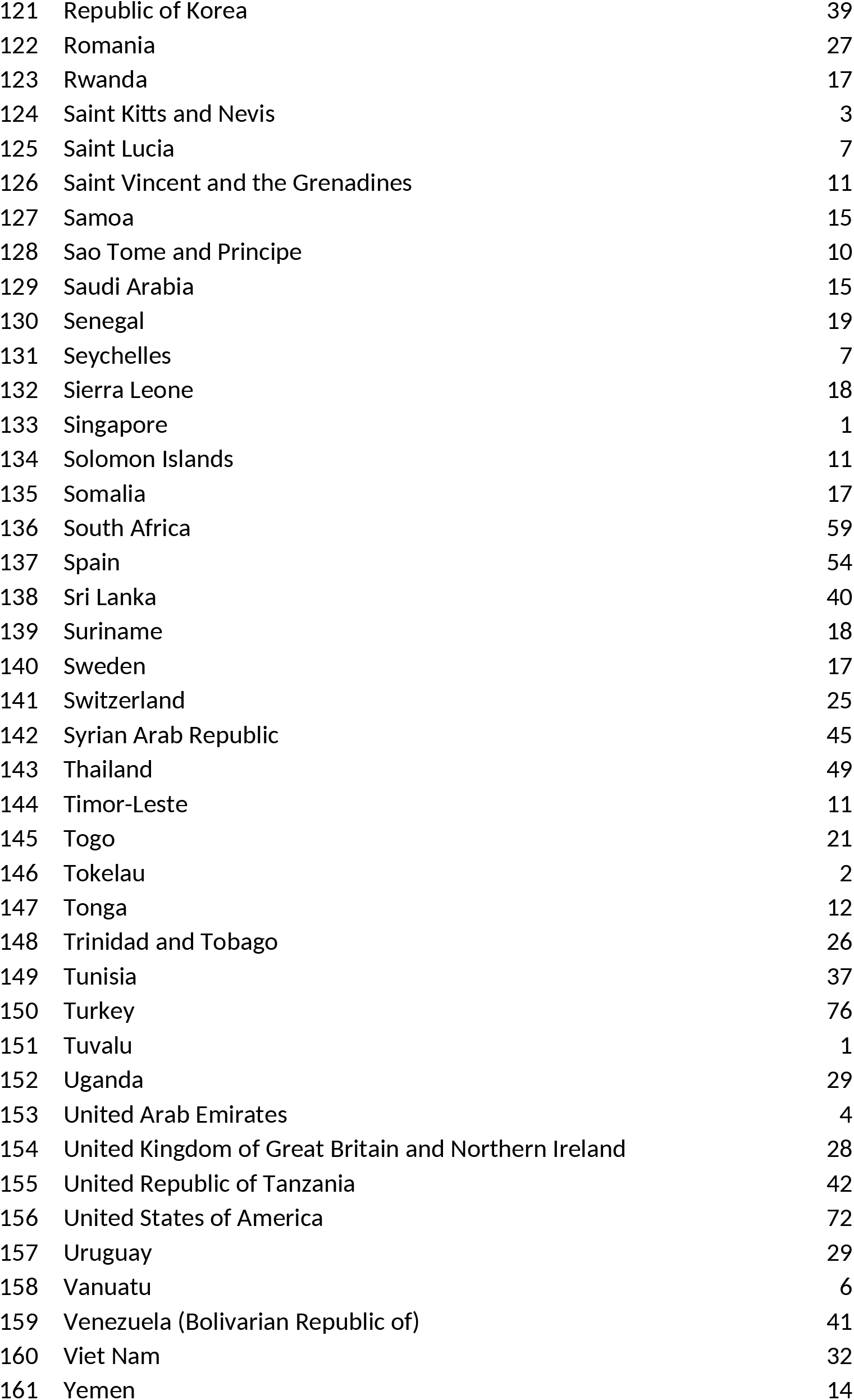
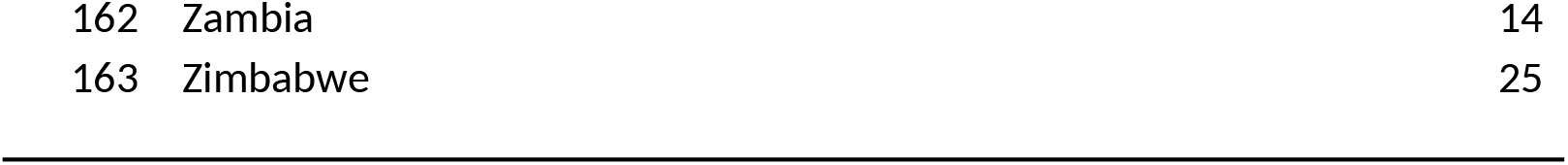
List of countries and number of crops per country with uninterrupted yearly yield records (1961-2020).

|  | **Country** | **Number of crops** |
| --- | --- | --- |
| 1 | Afghanistan | 23 |
| 2 | Albania | 15 |
| 3 | Algeria | 35 |
| 4 | Angola | 25 |
| 5 | Antigua and Barbuda | 12 |
| 6 | Argentina | 46 |
| 7 | Australia | 60 |
| 8 | Austria | 26 |
| 9 | Bahamas | 7 |
| 10 | Bahrain | 6 |
| 11 | Bangladesh | 47 |
| 12 | Barbados | 12 |
| 13 | Belize | 8 |
| 14 | Benin | 28 |
| 15 | Bhutan | 11 |
| 16 | Bolivia (Plurinational State of) | 46 |
| 17 | Botswana | 7 |
| 18 | Brazil | 52 |
| 19 | Brunei Darussalam | 10 |
| 20 | Bulgaria | 36 |
| 21 | Burkina Faso | 18 |
| 22 | Burundi | 19 |
| 23 | Cabo Verde | 11 |
| 24 | Cambodia | 18 |
| 25 | Cameroon | 30 |
| 26 | Canada | 41 |
| 27 | Central African Republic | 21 |
| 28 | Chad | 16 |
| 29 | Chile | 47 |
| 30 | China | 68 |
| 31 | China, Taiwan Province of | 49 |
| 32 | Colombia | 40 |
| 33 | Comoros | 8 |
| 34 | Congo | 23 |
| 35 | Cook Islands | 10 |
| 36 | Costa Rica | 23 |
| 37 | Côte d'Ivoire | 32 |
| 38 | Cuba | 28 |
| 39 | Cyprus | 38 |

**Table S2** (cont.)
|  |  |  |
| --- | --- | --- |
| 40 | Democratic People's Republic of Korea | 29 |
| 41 | Democratic Republic of the Congo | 46 |
| 42 | Denmark | 16 |
| 43 | Dominica | 10 |
| 44 | Dominican Republic | 34 |
| 45 | Ecuador | 56 |
| 46 | Egypt | 56 |
| 47 | El Salvador | 27 |
| 48 | Equatorial Guinea | 7 |
| 49 | Eswatini | 16 |
| 50 | Faroe Islands | 1 |
| 51 | Fiji | 19 |
| 52 | Finland | 20 |
| 53 | France | 53 |
| 54 | French Polynesia | 21 |
| 55 | Gabon | 13 |
| 56 | Gambia | 7 |
| 57 | Germany | 24 |
| 58 | Ghana | 29 |
| 59 | Greece | 47 |
| 60 | Grenada | 13 |
| 61 | Guatemala | 30 |
| 62 | Guinea | 22 |
| 63 | Guinea-Bissau | 11 |
| 64 | Guyana | 14 |
| 65 | Haiti | 29 |
| 66 | Honduras | 29 |
| 67 | Hungary | 32 |
| 68 | Iceland | 6 |
| 69 | India | 67 |
| 70 | Indonesia | 44 |
| 71 | Iran (Islamic Republic of) | 47 |
| 72 | Iraq | 34 |
| 73 | Ireland | 12 |
| 74 | Israel | 49 |
| 75 | Italy | 49 |
| 76 | Jamaica | 39 |
| 77 | Japan | 53 |
| 78 | Jordan | 32 |
| 79 | Kenya | 39 |
| 80 | Kiribati | 3 |

**Table S2** (cont.)
|  |  |  |
| --- | --- | --- |
| 81 | Kuwait | 3 |
| 82 | Lao People's Democratic Republic | 23 |
| 83 | Lebanon | 47 |
| 84 | Lesotho | 8 |
| 85 | Liberia | 17 |
| 86 | Libya | 28 |
| 87 | Madagascar | 51 |
| 88 | Malawi | 24 |
| 89 | Malaysia | 32 |
| 90 | Maldives | 8 |
| 91 | Mali | 24 |
| 92 | Malta | 2 |
| 93 | Mauritania | 13 |
| 94 | Mauritius | 24 |
| 95 | Mexico | 70 |
| 96 | Mongolia | 4 |
| 97 | Morocco | 47 |
| 98 | Mozambique | 32 |
| 99 | Myanmar | 32 |
| 100 | Namibia | 4 |
| 101 | Nauru | 1 |
| 102 | Nepal | 16 |
| 103 | Netherlands | 21 |
| 104 | New Caledonia | 9 |
| 105 | New Zealand | 38 |
| 106 | Nicaragua | 20 |
| 107 | Niger | 17 |
| 108 | Nigeria | 40 |
| 109 | Niue | 6 |
| 110 | Norway | 16 |
| 111 | Oman | 6 |
| 112 | Pakistan | 42 |
| 113 | Panama | 23 |
| 114 | Papua New Guinea | 22 |
| 115 | Paraguay | 29 |
| 116 | Peru | 67 |
| 117 | Philippines | 42 |
| 118 | Poland | 22 |
| 119 | Portugal | 38 |
| 120 | Puerto Rico | 19 |

**Table S2** (cont.)
|  |  |  |
| --- | --- | --- |
| 121 | Republic of Korea | 39 |
| 122 | Romania | 27 |
| 123 | Rwanda | 17 |
| 124 | Saint Kitts and Nevis | 3 |
| 125 | Saint Lucia | 7 |
| 126 | Saint Vincent and the Grenadines | 11 |
| 127 | Samoa | 15 |
| 128 | Sao Tome and Principe | 10 |
| 129 | Saudi Arabia | 15 |
| 130 | Senegal | 19 |
| 131 | Seychelles | 7 |
| 132 | Sierra Leone | 18 |
| 133 | Singapore | 1 |
| 134 | Solomon Islands | 11 |
| 135 | Somalia | 17 |
| 136 | South Africa | 59 |
| 137 | Spain | 54 |
| 138 | Sri Lanka | 40 |
| 139 | Suriname | 18 |
| 140 | Sweden | 17 |
| 141 | Switzerland | 25 |
| 142 | Syrian Arab Republic | 45 |
| 143 | Thailand | 49 |
| 144 | Timor-Leste | 11 |
| 145 | Togo | 21 |
| 146 | Tokelau | 2 |
| 147 | Tonga | 12 |
| 148 | Trinidad and Tobago | 26 |
| 149 | Tunisia | 37 |
| 150 | Turkey | 76 |
| 151 | Tuvalu | 1 |
| 152 | Uganda | 29 |
| 153 | United Arab Emirates | 4 |
| 154 | United Kingdom of Great Britain and Northern Ireland | 28 |
| 155 | United Republic of Tanzania | 42 |
| 156 | United States of America | 72 |
| 157 | Uruguay | 29 |
| 158 | Vanuatu | 6 |
| 159 | Venezuela (Bolivarian Republic of) | 41 |
| 160 | Viet Nam | 32 |
| 161 | Yemen | 14 |

**Table S2** (cont.)
|  |  |  |
| --- | --- | --- |
| 162 | Zambia | 14 |
| 163 | Zimbabwe | 25 |

**Table S3.** Summary of statistical results associated with the effects of the focal factors pollinator dependence and growth form on the probability of yield decline (*i.e.*, the probability that a given crop in a given country shows an average annual growth rate in yield <0 over a period of time), as evaluated separately by models GLMM_1a, GLMM_1b, respectively, and jointly by GLMM_2. The comparisons involve (1) the estimation of yield decline and their analyses considering the entire period 1961-2020, as it was carried in this study and reported in Table 1, *vs*. considering the period 1962-2019, thus curtailing the modal and expected highly influential years 1961 and 2020 (Fig. 1); and (2) considering pollinator dependence as a categorical variable with three levels (*i.e*., none, moderate, and high), as it was carried out in this study after lumping, *vs*. considering pollinator dependence as a numerical variable with five possible values (*i.e.*, 0, 5, 25, 65, 95% corresponding to the categories none, little, modest, high and essential, respectively), as analyzed in Deguines et al. (2014). As expected, the estimate of the slope of the probability of yield decline with increasing pollinator dependence, when this variable was treated as numerical, was positive (logit estimate +1SE = 0.005478 + 0.002203)

| Time series:<br>Pollinator dependence: | 1961-2020<br>Categorical |  |  | 1962-2019<br>Categorical |  |  | 1961-2020<br>Numerical |  |  |
| --- | --- | --- | --- | --- | --- | --- | --- | --- | --- |
| | Df | $\chi^2$ | P | Df | $\chi^2$ | P | Df | $\chi^2$ | P |
| **Model/ Focal factor** |  |  |  |  |  |  |  |  |  |
| GLMM_1a |  |  |  |  |  |  |  |  |  |
| Pollinator dependence | 2 | **6.71** | 0.035 | 2 | 5.55 | 0.062 | 1 | **6.14** | 0.013 |
| GLMM_1b |  |  |  |  |  |  |  |  |  |
| Growth form | 2 | **15.95** | 0.00034 | 2 | **16.08** | 0.00032 | 2 | **15.95** | 0.00034 |
| GLMM_2 |  |  |  |  |  |  |  |  |  |
| Pollinator dependence | 2 | 2.08 | 0.35 | 2 | 1.57 | 0.46 | 1 | 1.54 | 0.21 |
| Growth form | 2 | **11.93** | 0.026 | 2 | **11.85** | 0.0027 | 2 | **11.04** | 0.004 |

